# Chronic Activity-Based Anorexia triggers a glial response in the hippocampus independent of intestinal epithelial Toll-Like Receptor 4

**DOI:** 10.64898/2026.01.29.702534

**Authors:** Léna Rousseau, Tanguy Demangeat, Colin Salaün, Clara Queguiner, Charlène Guérin, Christine Bôle-Feysot, Oudiayé Maiga, Adam Tiffay, Fatima Léon, Léa Cornaille, David Ribet, Jean-Claude do Rego, Jean-Luc do Rego, Ludovic Langlois, Moïse Coëffier

## Abstract

Anorexia nervosa is characterized by maladaptive eating behavior and cognitive dysfunction, which could be explained by a neuroinflammation. A gut dysbiosis could link gastrointestinal alterations to central dysfunctions, particularly via the toll-like receptor 4 (TLR4), which has been shown to play a key role in the activity-based anorexia (ABA) model. We aimed to evaluate the neuroinflammation and its behavioral consequences in the ABA model, and to decipher the role of the microbiota-gut-brain axis, and more specifically of TLR4, in these alterations of the central nervous system. We show that chronic restriction is more strongly associated with gut inflammation, cecal microbiota alteration and neuroinflammatory processes in the hippocampus than acute restriction. The hippocampal glial response is characterized by a loss of astrocyte density, and an increased number of deramified microglia. We further demonstrate that these alterations are independent of TLR4 expressed by intestinal epithelial cells. In conclusion, our results highlight that the chronicity of ABA-associated undernutrition alters the response of glial cells in the hippocampus that is linked with changes in microbiota composition, highlighting the importance of faster diagnosis and treatment of AN.

## Introduction

Anorexia nervosa (AN) is an eating disorder characterized by a severe body weight loss, leading to a body mass index below 18.5 kg/m^2^. This condition arises from self-starvation and other restrictive behaviors such as problematic physical activity, misuse of laxatives or purging. A disturbed self-body image perception, known as dysmorphophobia, and an intense fear of gaining weight lead to the maintenance and worsening of this maladaptative eating behavior(1). The lifetime prevalence of AN is 1.4% in women and 0.2% in men and has dramatically increased in the last decades(2). Numerous comorbidities have been described in AN, including psychiatric comorbidities such as anxiety disorders and depression, but also somatic complications such as gastrointestinal disorders(3). AN is recognized as a multifactorial disease influenced by biological, genetic, environmental and psychosocial factors(4). Despite this, there is still no specific and effective treatment, with current care focusing on refeeding, prescription of antidepressants and psychological therapies(3). Thus, a better understanding of underlying mechanisms is needed and requires the use of preclinical models. The most commonly used approach is the activity-based anorexia (ABA) model, which combines food restriction and a free access to a running wheel. This model summarizes many features of AN, including sexual dimorphism, behavioral changes and dysfunction of the microbiota-gut-brain axis(5).

The microbiota-gut-brain axis appears to play a key role in AN, potentially linking gastrointestinal alterations to the maladaptive eating behavior and cognitive dysfunction observed in this pathology. A peripheral, chronic, low-grade inflammation has been described in AN(6),(7) and ABA model(8), associated with an altered intestinal barrier function(9),(10). The involvement of the gut microbiota is strongly suggested, as a dysbiosis is reported in mouse model(11) and in patients with AN(12), knowing that microorganisms of the gut lumen are major contributors to the integrity of the intestinal barrier(13). In addition, fecal microbiota transplantation from AN patients to germ-free mice subjected to calorie restriction induced reduced weight gain and an altered hypothalamic expression of energy metabolism and eating behavior markers(12). Toll-like receptor 4 (TLR4), a key player in microbiota-host communication that recognizes bacterial lipopolysaccharides and triggers pro-inflammatory signaling, seems to be of particular interest. Indeed, it is upregulated in intestinal epithelial cells and macrophages in ABA mice and its expression level correlates with cytokine production(8). Finally TLR4 deletion was shown to modified the ABA model response(14).

Proinflammatory cytokines are able to cross(15) and/or to increase the permeability(16) of the blood-brain barrier, reaching the brain and triggering a central inflammation called neuroinflammation and altering the hypothalamic energy balance(15). Neuroinflammation involves several mechanisms and actors including microglial and astrocytic cells. Through the mobility of their processes and the synthesis of inflammatory factors, glial cells act to initiate, regulate and inhibit inflammatory responses to protect the brain. The existence of a neuroinflammation in AN has been suggested, particularly in brain regions involved in the regulation of eating behavior, both in its homeostatic and hedonic dimensions. Several findings support this hypothesis, such as the reduced density of astrocytes in the hippocampus(17), prefrontal cortex and corpus callosum(18) in a dehydration-induced anorexia rat model. These changes were correlated with a lower brain volume, a feature also found in patients with AN(19,20). In addition, this rat model induced a deramification of astrocytes(21) and microglia(22), as well as an enhanced expression of proinflammatory cytokines. Thus, neuroinflammation could participate in the core features of AN and, the non-physiological response to starvation, but also in some of the comorbidities associated with AN, such as anxiety and cognitive disorders.

The aim of this work was to evaluate the neuroinflammation and its behavioral consequences in the ABA model at the acute and chronic phases, and to decipher the link with alterations of the microbiota-gut-brain axis, and more specifically with the intestinal TLR4.

## Materials and methods

### 1. Animal experiments

#### Experimental procedure

6-week-old females C57BL/6JRj mice were purchased from Janvier Labs (Le Genest St Isle, France). After 5 days of acclimatization to inversed cycle (dark phase between 10:00 AM and 10:00 PM) and a standard diet (TEKLAD Global Diets 2918, Inotiv, US) at 4 mice per cages in the SCAC behavioral analysis platform (UFR Santé, Rouen, France), mice were randomized according to body weight in either the control (CT), limited food-access (LFA) or activity-based anorexia (ABA) groups (n=18/group for acute model, n=24/group for chronic model), and housed 1 per cage. CT and LFA mice were placed in standard cages and ABA mice in cages equipped with a monitored running wheel (ActiviWheel software Intellibio, Seichamps, France). Food intake, water intake, body weight and welfare scoring were measured daily at the end of the light phase. All mice were fed *ad libitum* for 1 week before starting the protocol of restriction for LFA and ABA mice adapted from previous studies(23–25). Food intake for the last 5 days of the habituation period was averaged for each LFA and ABA mouse to establish a reference value of food intake. A quantitative restriction of 60% of the individual reference value was then applied for 1 week, to reach a 25% body weight loss, constituting the acute phase of the restriction. This target weight loss was then maintained until 17 days (acute model) or until 43 days (chronic model), with a 50% food restriction adjusted each day according to the change in daily body weight. The restriction was adjusted by plus or minus 5% for every 2.5% increment in weight loss deviating from the target weight. Food access was provided at the beginning of the dark phase (10:00 AM) and placed directly on the ABA and LFA’s bedding. Experimental procedure was approved by the local ethical committee and the ministerial committee for animal experimentation (#47092-2024012616077845).

#### Genetic model

CreVillin^ERT2^ (gift from Dr Sylvie Robine, Institut Curie, Paris, France) and TLR4 mice (Jackson Laboratory, US) were crossed to obtain CreVillin^ERT2^ TLR4 mice. Genotype was checked *post-mortem* as previously described(14). Mice were housed at the University of Rouen main animal facility (UFR Santé, Rouen, France) during all the breeding process. 72 females 5-week-old CreVillin^ERT2^ ^+/+^ TLR4^floxed^ ^+/+^ mice were transferred to the SCAC behavioral analysis platform (UFR Santé, Rouen, France) for the acclimatization to inversed cycle (dark phase between 10:00 AM and 10:00 PM) and a standard diet (TEKLAD Global Diets 2918) at 4 mice per cages, for 5 days. Mice were randomized according to body weight in either the invalidated (TLR4^IEC-/-^, n=36) or non-invalidated group (TLR4^IEC+/+^, n=36) and were submitted to 5 days of intraperitoneal (i.p.) injections of 100 μL tamoxifen (T5648, Sigma-Aldrich, Saint-Louis, US) at 10 mg/mL in 10% ethanol and 90% sunflower oil as previously described(26), in order to invalidate TLR4 specifically in intestinal epithelial cells (IEC). A second randomization according to body weight was performed to constitute 6 groups: control (CT_TLR4^IEC+/+^ and CT_ TLR4^IEC-/-^), limited food-access (LFA_TLR4^IEC+/+^ and LFA_ TLR4^IEC-/-^) or activity-based anorexia (ABA_TLR4^IEC+/+^ and ABA_TLR4^IEC-/-^) groups (n=12/group), housed 1 per cage. These mice were then subjected to the same chronic experimental procedure as described above.

#### Nesting test

Nestlet shredding test and nest building test were carried out as previously described(27). Briefly, environmental enrichments were removed, except for the running wheel, from the cage 24h prior to the first test day. Housing cages of the mice were used as test areas.

Nestlets were handled with forceps, weighted and placed in the cage. After 24h, before the daily handling of mice, 3 independent investigators that were not involved in the testing procedure graded nests using published scores(28): from 1 (very poor/no nest building) to 5 (optimal nest building). Then, the largest part of the nestlets was removed and weighted to calculate the percent of shredded material.

#### Light-dark box test

Anxiety and exploratory behaviors in response to an aversive environment were assessed using the light-dark box test. The system is made of a bright (200 lux) compartment (1/2) and a dark compartment (1/2) connected by an open door. In the nocturnal phase, mice were placed 15 min in the light in the experimental room, then positioned in the light compartment, next to the door for 15 min. We assessed the time and distance travelled in each compartment, and infra-red lasers allowed horizontal and vertical movements to be distinguished. Recording was performed by Fusion 6.4 software (Omnitech Electronics Inc., Columbus, Ohio, US).

#### Tissue collection

At the end of the experiment, at 10:00 AM, mice were anesthetized and analgesized with an i.p. injection of ketamine (100 mg/kg, Centravet, Plancoet, France) and xylazine (10 mg/kg, Bayer Healthcare, Puteaux, France) diluted in NaCl 0.85%, buprenorphine (0,05 mg/kg, Buprecare, Coveto, France). Blood was collected from the caudal *vena cava* on heparinized vacutainer tubes, centrifuged at 3000 g for 15 min at 4°C and then plasma was collected. All samples (plasma, hypothalamus, hippocampus, amygdala, ileum, colon and cecal content) were immediately frozen in liquid nitrogen and stored at −80°C until analysis. Jejunal fragments were prepared following Swiss-rolling technique as previously described(29). Half of the whole brains and jejunal wraps were immediately fixed by immersion in paraformaldéhyde (4% in PBS, Thermo fisher Scientific, Waltham, MA, US) for 24 hours, then transferred to D-sucrose (30%, VWR Chemicals, Rosny-sous-bois, France) for 72 hours at 4°C. Tissues were then frozen in isopentane (VWR Chemicals) at −30°C, then stored at −80°C.

### 2. Ex vivo colonic permeability by Ussing chambers

Colonic permeability was assessed by measuring fluorescent molecules (Lucifer Yellow, 400 Da, Merck, Germany and Dextran Alexa Fluor 680, 3 kDa, Thermo Fisher Scientific, US) mucosal to serosal flux level in Ussing chambers (Harvard Bioscience, Multi Channel Systems MCS GmbH, Reutlingen, Germany) as previously described(30). Samples were maintained at 37°C during 3h, then serosal medium was collected, and fluorescence levels were measured with a Spark multimode microplate reader (Tecan, Männedorf, Switzerland). In addition, trans-epithelial electric resistance (TEER) was measured.

### 3. Immunofluorescence Free-floating sections

Fixed frozen brains were cut in serial coronal 40 μm-thick sections using a CM 1950 cryostat (Leica Biosystems, Nussloch, Germany), within the following Bregma points: 1.54 to 0.74 mm for nucleus accumbens, −0.70 to −1.06 mm for paraventricular nucleus, −1.22 to −1.58 mm for arcuate nucleus and −1.46 to −1.94 mm for dentate gyrus and CA3, based on the mouse brain atlas (The Mouse Brain in Stereotaxic Coordinates, 3^rd^ Edition, Franklin and Paxinos, 2008).

Free-floating sections protocol was adapted from previous data(31). Brain sections were washed three times in PBS buffer before antigen retrieval for 30 min in a 10 mM sodium citrate solution at pH 8.5-9.0 preheated and maintained at 80°C in a water-bath as recommended(32). Sections were rehydrated for 30 min in PBS, and then placed in blocking solution (10% bovine serum albumin, 0.3% Triton X-100 in PBS) for 30 min. Sections were incubated overnight at 4°C in a shaker with the following primary antibodies: rat anti-GFAP monoclonal antibody (1/1000e, 13-0300, Invitrogen, Carlsbad, CA, US), and rabbit anti-IBA1 polyclonal antibody (1/2000e, 01919741, Wako, Neuss, Germany) in the incubation solution (1% bovine serum albumin, 0.1% Triton X-100 in PBS). After three washing steps, sections were exposed to the appropriate secondary antibody for 2h in the dark at room temperature: AlexaFluor 488 goat anti-rat antibody (1/400e, AlexaFluor A11008, Life Technologies-Invitrogen) and AlexaFluor 555 goat anti-rabbit antibody (1/400e, AlexaFluor A21428, Life Technologies-Invitrogen). This step was followed by washing and mounting on Superfrost Plus microscope slides (Epredia, Breda, Netherlands). Sections were counterstained with Fluoroshield with DAPI (Sigma-Aldrich), covered by coverslip (Epredia) and stored at 4°C until microscopic observation. Nonspecific coupling of secondary antibodies was checked by incubation of sections without primary antibodies.

#### Image acquisition

##### Fluorescent microscopic acquisition for density analysis

Images (2048×2048 pixels, pixel size 0.33 μm) were obtained using a widefield fluorescence Leica Thunder Imager Tissue 3D microscope (Leica Biosystems) and Las X Navigator software (Leica Microsystems, Wetzlar, Germany). Left and right bilateral structures of 2 successive sections per region per mice were acquired in a blind-fashion. DAPI, GFAP and IBA1 fluorescence was detected with filter cubes 390, GFP and Y3, respectively. Z-stacks (intervals of 0.5 μm) were taken with a DFC9000 GT VSC-12292 monochrome camera and HC PL APO objective at x20 magnification with identical illumination and exposure settings for all recording.

##### Confocal acquisition for morphology analysis

Images (1024×1024 pixels, pixel size 0.23 μm) of dentate gyrus were obtained using a confocal TCS SP8 DM6000B-CFS microscope (Leica Biosystems) and Las X Navigator software (Leica Microsystems), as previously described(33). 1 picture per hemisphere per mice was acquired in a blinded-fashion, using the mosaic system. GFAP and IBA1 fluorescence were located with hybrid detectors. Sequential z-stacks (intervals of 0.3 μm) were taken with a Leica DFC 365 Fx camera at x63 magnification (oil-immersion lens, zoom 0.75) with identical confocal settings.

#### Analysis

##### Density analysis

Widefield fluorescence images were used to study density of GFAP^+^ and IBA1^+^ cells in each region of interest (ROI): arcuate nucleus (ARC), paraventricular nucleus (PVN), dentate gyrus (DG), *cornu ammonis 3* (CA3) and accumbens nucleus (ACC) (2 sections per hemispheres per brain region per mice), using ImageJ software (National Institutes of Health, Bethesda, Maryland, US). Images treatment was done automatically with a macro designed for this purpose (sum projection of z-stacks, area measure of the ROI, DAPI counting). All cells in the section plane were counted and reported as number of cells per 0.1 mm^2^.

##### Morphological analysis

Confocal images were analyzed to study morphological parameters of GFAP^+^ and IBA1^+^ cells using IMARIS software (version 10.1.0, Oxford Instruments), as previously described(33). 2 to 4 cells per hemisphere per mouse were selected if their soma was present in the section plane and their extensions were not cut by the x and/or y axis. The 3D reconstruction of IBA1^+^ cells was carried out with the ‘Surface’ tool indicating the volume of the cell. The extensions of IBA1^+^ and GFAP^+^ cells as well as the soma of IBA1^+^ cells were traced semi-automatically using the ‘Filaments’ tool. Thus, filament length, full branch level and depth, number of terminal and branch points and Sholl analysis were obtained. IBA1^+^ cells soma parameters, like volume, area, prolate ellipticity and ellipsoid axis higher length (C) were calculated with the ‘Filaments’ tools.

### 4. RNA extraction and reverse transcription

TRIzol-chloroform (Life Technologies-Invitrogen and Sigma-Aldrich, Darmstadt, Germany, respectively) RNA extractions were performed as previously described(34) on hypothalamus, hippocampus, amygdala, ileum and colon. RNAs quality and quantity were assessed using Nanodrop® (Thermo fisher Scientific, Waltham, MA, US). 1 μg of RNAs was treated with DNAse (Promega, Charbonnières-les-Bains, France) and used for reverse transcription into cDNAs with M-MLV (Life Technologies-Invitrogen).

### 5. Real-time quantitative polymerase chain reaction (RT-qPCR)

RT-qPCR were performed with SYBR Green technology (Bio-Rad Laboratories, Marnes la Coquette, France) on CFX 96 Real-Time PCR System (Bio-Rad, Hercules, CA, US). Primers sequences are listed in Additional file 1. The values were obtained by the conversion of cycle thresholds to concentration values using standard curves. mRNA levels of genes of interest were normalized by the mean of mRNA levels of reference genes (*Rps18* and *Eef2*).

### 6. Proteins extraction

Colon proteins were extracted using 2 different lysis buffers adapted to the analysis of the different protein targets studied: buffer A (Hepes pH 7.9 10 mM, KCl 10 mM, MgCl2 1.5 mM, EDTA 0.1 mM), DTT (1 mM), NP40 (0.25%), protease (0.5%) and phosphatase (1%) inhibitors) was used for the study of claudin-1; buffer U-CHAPS (spermine tetrahydrochloride 25 mM, protease (5%) and phosphatase (1%) inhibitors (Sigma-Aldrich), DTT 50 mM (Janssen Pharmaceuticalaan, Geel, Belgium), Urea 8 M (Merck Millipore, Burlington, US), CHAPS 4% (Janssen Pharmaceuticalaan)) for the study of occludin and ZO-1. Total protein content was determined using Bradford assay and 2D Quant kit (GE Healthcare, Buc, France), respectively.

### 7. Evaluation of protein expression by western blot

Total proteins (20 μg/lane) were separated on 4-20% (claudin-1) or 10% (occludin and ZO-1) gradient polyacrylamide stain-free gel (Bio-Rad Laboratories). After electrophoresis (100 V for 1h), proteins were transferred onto nitrocellulose (Bio-Rad Laboratories) or PVDF membranes for claudin-1 or occludin and ZO1, respectively. After blockage for 1h in milk 5%, membranes were incubated overnight at 4°C with primary antibody: anti-claudin-1 (71-7800, Thermo fisher Scientific, Waltham, MA, US), anti-occludin (OC-3F10, Thermo fisher Scientific) and anti-ZO-1 (40-2200, Thermo fisher Scientific) at 1/1000^e^. Membranes were washed before adequate secondary antibody incubation for 1h (Agilent Technologies Denmark, Glostrup, Denmark, 1/5000^e^) at room temperature. Immunocomplexes were revealed using a chemiluminescence detection system (Clarity^TM^ Western ECL substrate, Bio-Rad Laboratories). Acquisition was performed on ChemiDoc MP Imaging System (Bio-Rad Laboratories). Results were analyzed on Image Lab^TM^ software (Bio-Rad Laboratories) and normalized to total proteins.

### 8. Jejunal histology

Swiss rolls were cut in 12 μm sections using a CM 1950 cryostat (Leica Biosystems) and stained with a solution of hematoxylin and eosin. Images (5472x3648 pixels, pixel size 0.24 μm) were obtained using a widefield Leica Thunder Imager Tissue 3D microscope (Leica Biosystems) with a brightfield DMC 5400 camera and Las X Navigator software (Leica Microsystems). 10 regions per mice were acquired in a blind-fashion at 10x magnification. Jejunal villi height and thickness were measured with ImageJ software as previously described(35).

### 9. Cecal DNA extraction and analysis

DNA from cecal contents were extracted using QIAamp Fast DNA Stool mini kit (Qiagen, Hilden, Germany) as described previously(11). The integrity of extracted DNA was assessed by visualizing their migration patterns on 1% agarose gels. The University of Minnesota Genomics Center performed Element AVITI24 sequencing as follows: the V5-V6 region of the 16S rRNA gene was PCR-enriched using the primer pair V5F_Nextera (TCGTCGGCAGCGTCAGATGTGTATAAGAGACAGRGGATTAGATACCC) and V6R_Nextera (GTCTCGTGGGCTCGGAGATGTGTATAAGAGACAGCGACRRCCATGCANCACCT) and then underwent a library tailing PCR as previously described(36). The amplicons were purified, quantified and sequenced using an Element AVITI24 to produce 2 x 300 bp sequencing products. Then, we performed quality control of demultiplexed sequencing data with FastQC version 0.12.1. No samples were discarded. Primers and adapters were eliminated using Cutadapt version 5.1. Then, the obtained sequences were denoised and chimera detected through DADA2(37) plugin of QIIME2 version 2025.7. The amplicon sequence variants (ASV) were associated to taxonomic identities using a pre-trained Naïve Bayes classifier with the SILVA v138.2 database and the plugin classify-sklearn. Rarefaction curves were generated to assess sequencing depth, and the data were rarefied to a common sampling depth prior to diversity analyses (acute model: 22899, chronic model: 77618). Alpha and beta diversity metrics were calculated with the QIIME2 diversity plugin(38). Data were normalized on the total number of ASV for differential analysis.

### 10. Statistical analysis

Results were expressed as mean ± standard error to mean. Statistical analyses and graphs were performed with GraphPad Prism 9 software (GraphPad Software Inc., San diego, CA, US). Outliers were excluded based on the ROUT method with a Q coefficient equal to 1%, and normal distribution was assessed using Shapiro-Wilk test. Brown-Forsythe test was used for testing the assumption of equal variances. Data of acute and chronic models were analyzed with 1-way ANOVA or Kruskal-Wallis followed by Tukey or Dunn post-tests, respectively, and TLR4 KO model with 2-way ANOVA followed by Tukey post-tests.

Microscopy data were analyzed using nested ANOVA to account for replicates interdependence. Correlations were computed using Spearman test and Benjamini-Hochberg corrections were applied for multiple testing. A permutational multivariate analysis of variance (PERMANOVA) test was performed on the Bray-Curtis matrices using 999 random permutations and at a significance level of 0.05. Comparisons with *p*<0.05 were considered statistically significant and mentioned with the following symbols when *p*<0.05 and exact *p* values were written when a trend was observed (*p*<0.1): * p<0.05, ** p<0.01, *** p<0.001, ****p<0.0001.

## Results

### Acute and chronic ABA models induce a body weight loss dependent of wheel activity presence

In the acute model, cumulated food intake was higher in ABA mice in the habituation phase compared to CT and LFA (p<0.0001, Fig.1b), then CT mice became significantly higher than LFA and ABA during the restriction phase (p<0.0001). Initiation of food restriction led to body weight loss in ABA and LFA mice compared to CT from D8 (p<0.01, Fig.1c), until it reached the target of 25% body weight loss at D13 (Additional file 2c). Body weight loss was more pronounced in LFA mice compared to ABA mice at the end of the protocol (p<0.0001, Fig.1c) despite a less severe food restriction (p<0.0001, Fig.1d). Physical activity recorded in ABA group was mostly nocturnal during the habituation week (Additional file 2e). Total activity increased after the first day of food restriction until D11, then dropped sharply. A shift between nocturnal and diurnal activity was observed, mainly due to an increase of food-anticipatory physical activity (3 hours before food access) compared to the habituation phase (p<0.05, Fig.1e).

Similar patterns were retrieved in the chronic model regarding cumulated food intake, body weight and physical activity (Fig.2): a higher cumulated food intake was measured in ABA than in CT in the habituation phase (p<0.05, Fig.2b); the restriction phase was associated with a lower cumulated food intake in LFA and ABA mice (p<0.0001), resulting in a rapid body weight loss of 25% reached at D13 in both LFA and ABA (Additional file 2d) and a shift in the physical activity pattern was observed (Additional file 2f and e). No difference in body weight loss was observed between LFA and ABA at the end of the protocol (D45, ns, Fig.2c). In addition, maintaining the target body weight required adjustments to the amount of food delivered, which varied over time (p(Time)<0.0001, Fig.2d). In order to maintain weight loss, food restriction had to be increased, more markedly in ABA compared to LFA mice (p<0.0001, Fig.2d).

**Fig. 1:**
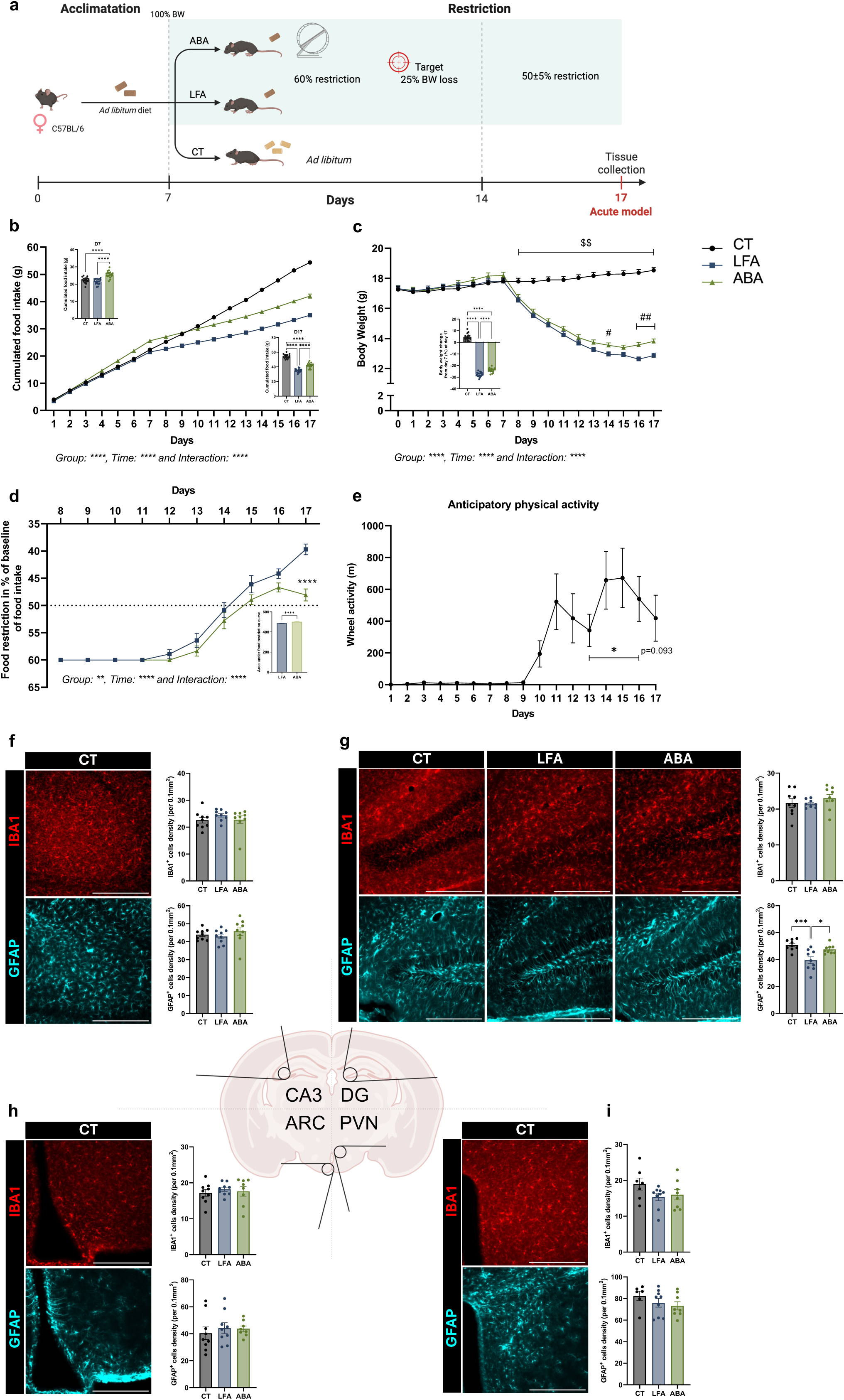
**Acute ABA model induces a wheel activity-dependent body weight loss and alters glial density.** Female C57BL/6J mice underwent a quantitative restriction of 60% of the individual reference value of food intake for 1 week, to reach a 25% body weight loss, then an adjusted 50% restriction to maintained it for 3 days (n=54). Mice had free access to a monitored running wheel (ABA; n=18) or not (limited-food access (LFA); n=18). Control mice were fed ad libitum and had no access to a running wheel (CT; n=18). **(a)** Experimental design of acute model. **(b)** Cumulated food intake (in g) during the protocol, at D7 and D17. **(c)** Body weight (in g) throughout the protocol and final body weight change from day 7 (in %). **(d)** Adjustments of food restriction in percentage of reference value of food intake, and area under the curve. **(e)** Wheel activity (in m) measured during 3 hours before food access. Ordinary one way ANOVA with Tukey’s multiple comparisons test for bar graphs, repeated measures two way ANOVA with Tukey’s multiple comparisons test for line graphs, mixed effects one way ANOVA with Dunnett’s multiple comparisons test with D7 for anticipatory physical activity graph. Widefield fluorescence acquisitions of IBA1^+^ (red) and GFAP^+^ (cyan) cells in the **(f)** CA3, **(g)** dentate gyrus (DG), **(h)** arcuate nucleus (ARC) and **(i)** paraventricular nucleus (PVN). Quantifications show the mean cell number per mouse (n=9 per group). Scale bar: 100 μm. Nested one-way ANOVA with Tukey’s multiple comparisons (4 to 8 replicas per mice). *p<0.05, **p<0.01, ***p<0.001, ****p<0.0001, $ CT vs ABA and LFA, # LFA vs ABA. Data are presented as mean values ^+^/- SEM.

**Fig. 2:**
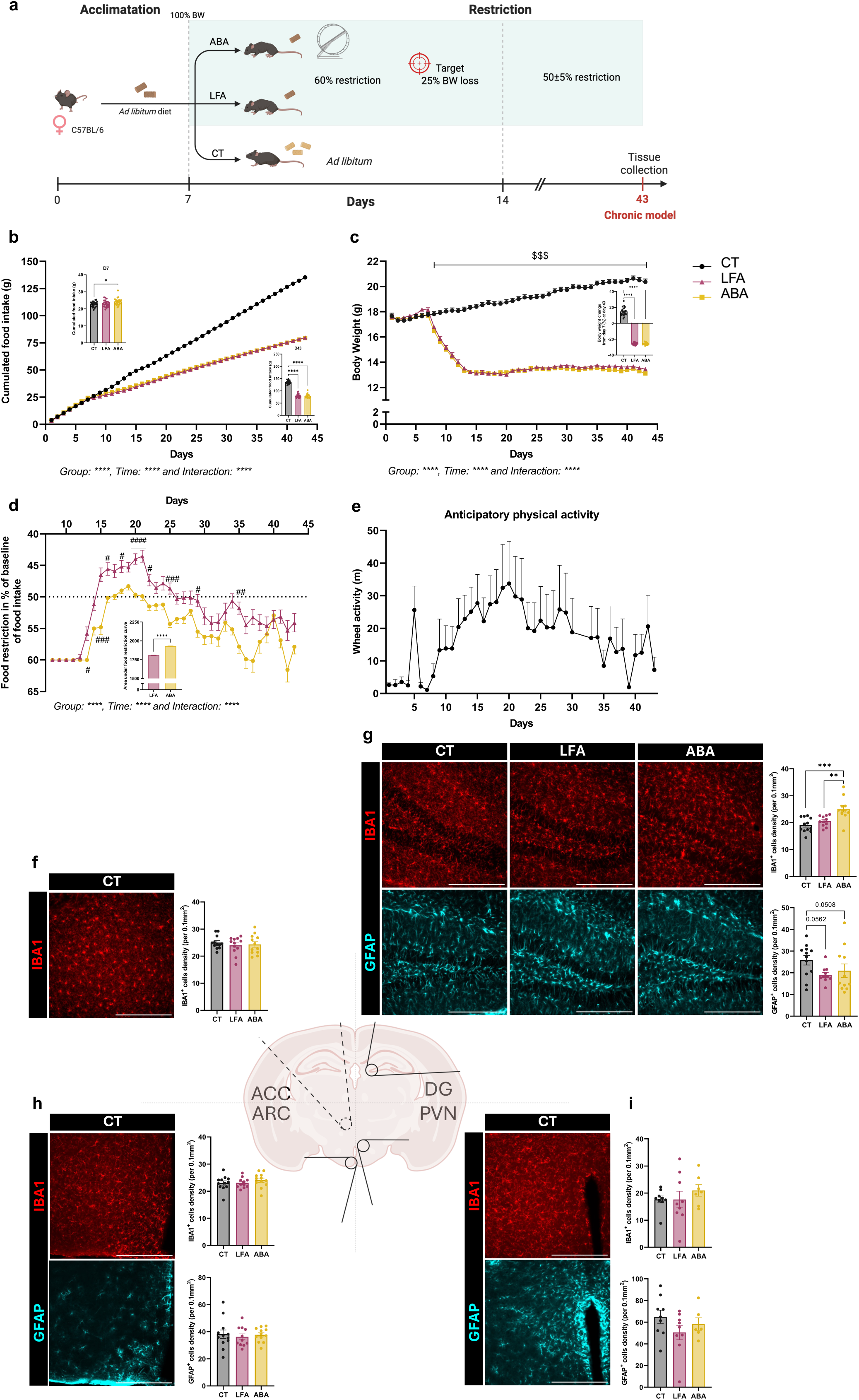
**Chronic ABA model induces a wheel activity-dependent body weight loss and alters glial density.** Female C57BL/6J mice underwent a quantitative restriction of 60% of the individual reference value of food intake for 1 week, to reach a 25% body weight loss, then an adjusted 50% restriction to maintained it until 43 days (n=36). Mice had free access to a monitored running wheel (ABA; n=24) or not (limited-food access (LFA); n=24). Control mice were fed ad libitum and had no access to a running wheel (CT; n=24). **(a)** Experimental design of chronic model. **(b)** Cumulated food intake (in g) during the protocol, at D7 and D43. **(c)** Body weight (in g) throughout the protocol and final body weight change from day 7 (in %). **(d)** Adjustments of food restriction in percentage of reference value of food intake, and area under the curve. **(e)** Wheel activity (in m) measured during 3 hours before food access. Ordinary one way ANOVA with Tukey’s multiple comparisons test for bar graphs, repeated measures two way ANOVA with Tukey’s multiple comparisons test for line graphs, mixed effects one way ANOVA with Dunnett’s multiple comparisons test with D7 for anticipatory physical activity graph. Widefield fluorescence acquisitions of IBA1^+^ (red) and GFAP^+^ (cyan) cells in the **(f)** accumbens nucleus (ACC), **(g)** dentate gyrus (DG), **(h)** arcuate nucleus (ARC) and **(i)** paraventricular nucleus (PVN). Quantifications show the mean cell number per mouse (n=12 per group). Scale bar: 100 μm. Nested one-way ANOVA with Tukey’s multiple comparisons (4 to 8 replicas per mice). *p<0.05, **p<0.01, ***p<0.001. Data are presented as mean values ^+^/- SEM.

### ABA model alters the glial density specifically in the DG

To decipher the glial response to ABA model, we carried out immunofluorescence staining of GFAP and IBA1 markers in brain regions involved in eating behavior. In the acute model, only the DG was impacted in terms of glial density, with a reduced GFAP^+^ cells number in LFA mice compared to CT and ABA (p<0.001 and p<0.05 respectively, Fig.1g). No difference was observed in the ARC, PVN or CA3 regions (ns, Fig.1f, h and i).

Similarly, the chronic model was associated, in the DG, with a slight drop of GFAP^+^ cells in restricted mice compared to CT (p=0.056 for LFA, p=0.051 for ABA, Fig.2g). In addition, the number of IBA1^+^ cells in the DG was increased in ABA mice (p<0.001 vs CT, p<0.01 vs LFA, Fig.2g).

### Glial cells adopt a specific phenotype in the DG in response to the chronic ABA model

As the chronic model seemed to alter glial density in the DG, we decided to investigate the three-dimensional morphometry of microglia and astrocytes in this region. Globally, a decrease of the studied parameters was reported in microglia of restricted mice (Fig.3). More specifically, total filament length, full branch level, number of terminal and branch points and total Sholl intersections were reduced in both LFA and ABA groups (Fig.3c). Full branch depth and maximal domain range were decreased in ABA (p<0.01) and LFA (p<0.05), respectively. A trend for a decrease in microglia volume was observed in ABA mice (p=0.055). We next performed Sholl analysis to quantify the ramification’s intersections with concentric spheres of 1 μm radius starting from the soma (Fig.3d). A strong reduction in the arborization pattern was observed in restricted mice (p(Group)<0.0001) and in a more pronounced manner in

**Fig. 3:**
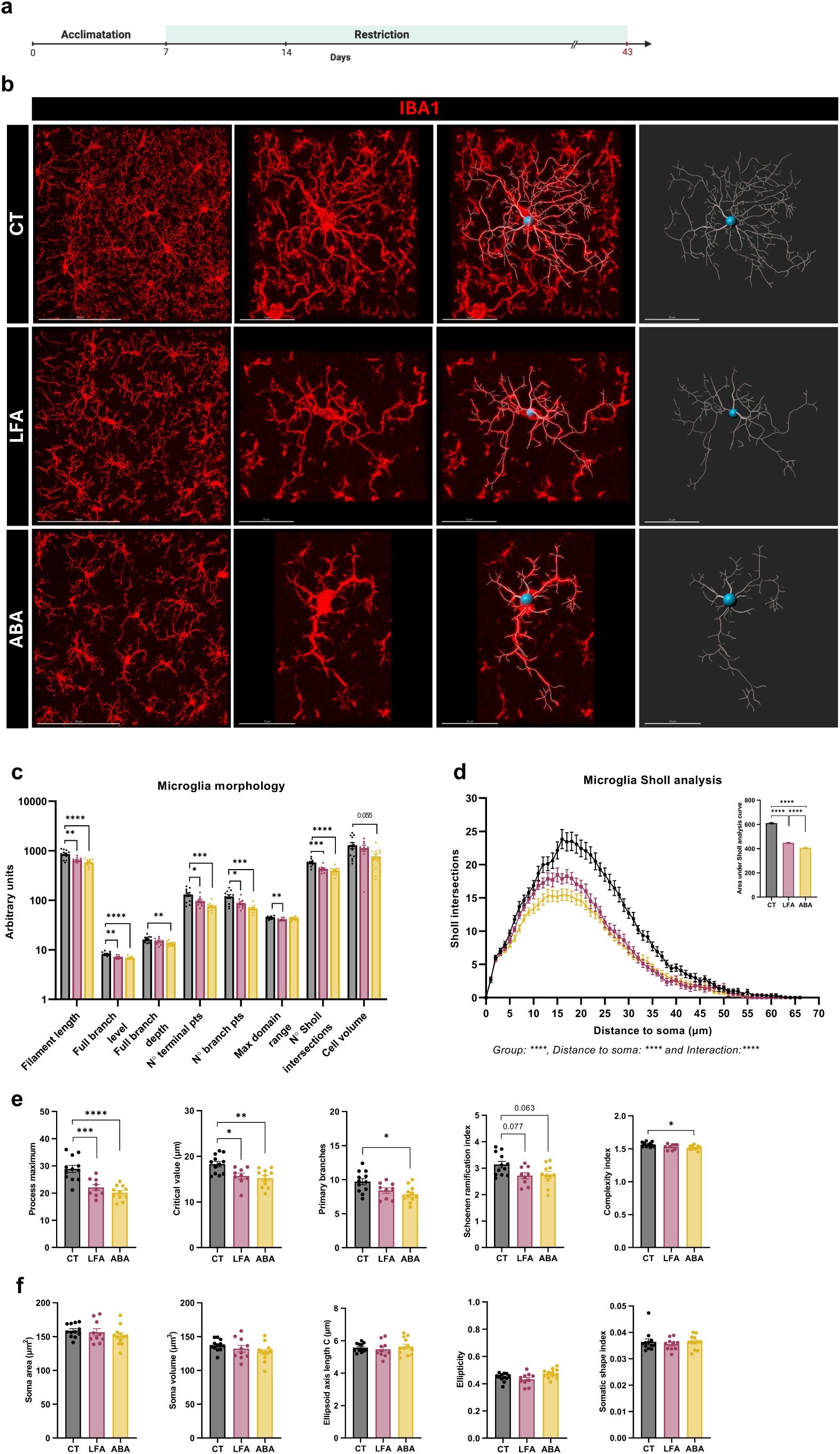
**Chronic ABA model induces a specific microglial phenotype in the DG.** Female C57BL/6J mice underwent a quantitative restriction of 60% of the individual reference value of food intake for 1 week, to reach a 25% body weight loss, then an adjusted 50% restriction to maintained it until 43 days (n=36). Mice had free access to a monitored running wheel (ABA; n=12) or not (limited-food access (LFA); n=12). Control mice were fed ad libitum and had no access to a running wheel (CT; n=12). **(a)** Experimental design of chronic model. **(b)** Confocal acquisitions of IBA1^+^ (red) cells in the dentate gyrus (DG) (scale bar: 100 μm), and zoom of a cell with morphometric reconstitution (scale bar: 50 μm). **(c)** Quantification of general morphometric parameters calculated by IMARIS software. **(d)** Number of microglial intersections over the distance to soma (Sholl analysis) and area under curve (AUC) **(e)** Quantification of Sholl parameters. **(f)** Quantification of soma morphological parameters. Quantifications show the mean value per mouse. Nested one-way ANOVA with Tukey’s multiple comparisons (3 to 4 cells per hemisphere per mice). *p<0.05, **p<0.01, ***p<0.001. Data are presented as mean values ^+^/- SEM.

ABA mice (AUC, p<0.0001 vs LFA, Fig.3d). Based on Sholl graph, we reported the following parameters as previously described(39): the highest number of intersections with Sholl spheres (process maximum) and the soma distance at which it occurs (critical value), the number of ramifications emerging from the soma (primary branches) and an index of overall branching (Schoenen ramification index, equal to process maximum/primary branches). A difference in number and distribution of processes was observed in restricted mice. Indeed, process maximum and critical value were lower in both LFA and ABA mice compared to CT (process maximum: p<0.001 and p<0.0001, critical value: p<0.05 and p<0.01, Fig.3e, respectively). Fewer primary branches were counted in ABA mice compared to CT (p<0.05, Fig.3e), leading to a trend for a decrease of Schoenen ramification index in restricted mice (p=0.077 for LFA and p=0.063 for ABA compared to CT, Fig.3e). Since the number of Sholl intersections did not directly reflect complexity and convolution degree(40), we evaluated the ratio between total process length and total intersection number, described as a complexity index. This ratio revealed a lower degree of complexity and convolution in ABA mice compared to CT (p<0.05, Fig.3e). As structural remodeling of microglia impacts not only branching but also soma size and shape, we focused on soma parameters(40). We did not highlight differences in soma area and volume, longest axis length or ellipticity (Fig.3f). The somatic shape index, that permits to distinguish round from rod-like microglia based on the deviation from circular shape did not show any differences between the three groups.

In astrocytes, branching parameters and Sholl analysis only revealed alterations in LFA mice (Fig.4). Indeed, full branch level and depth were decreased in LFA compared to ABA (p<0.05) and CT (p<0.05, Fig.4c). A group effect (p(Group)<0.01) was observed in Sholl analysis, with a decreased area under the curve in LFA mice (p<0.001 vs CT and p<0.01 vs ABA, Fig.4d). Sholl-derived parameters like process maximum, critical value, primary branches or Schoenen ramification index remained unchanged (Fig.4e). GFAP staining did not permit us to study soma morphometry.

**Fig. 4:**
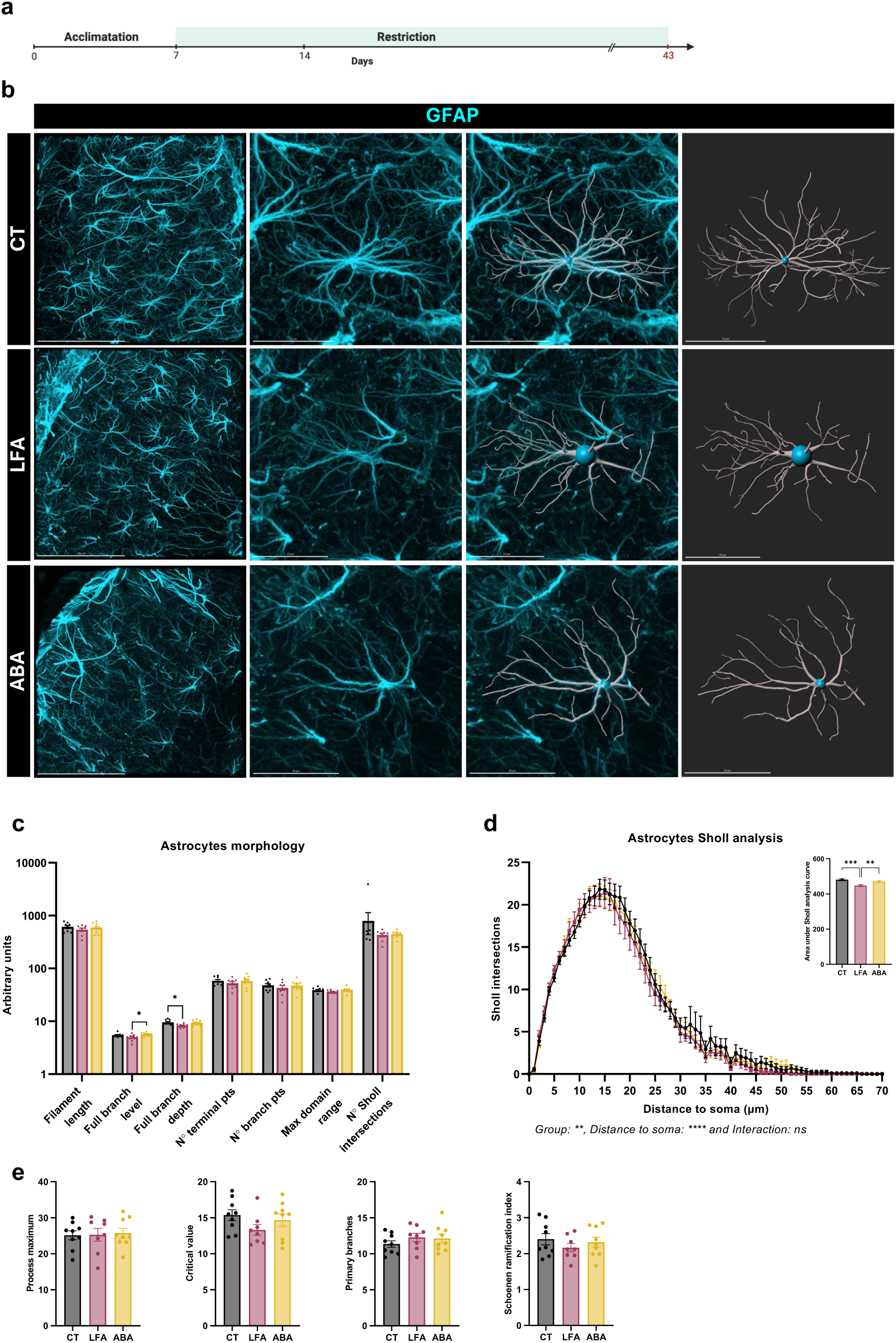
**Chronic ABA model induces a specific astrocytic phenotype in the DG.** Female C57BL/6J mice underwent a quantitative restriction of 60% of the individual reference value of food intake for 1 week, to reach a 25% body weight loss, then an adjusted 50% restriction to maintained it until 43 days (n=36). Mice had free access to a monitored running wheel (ABA; n=12) or not (limited-food access (LFA); n=12). Control mice were fed ad libitum and had no access to a running wheel (CT; n=12). **(a)** Experimental design of chronic model. **(b)** Confocal acquisitions of GFAP^+^ (cyan) cells in the dentate gyrus (DG) (scale bar: 100 μm), and zoom of a cell with morphometric reconstitution (scale bar: 25 μm). **(c)** Quantification of general morphometric parameters calculated by IMARIS software. **(d)** Number of astrocytic intersections over the distance to soma (Sholl analysis) and area under curve (AUC) **(e)** Quantification of Sholl parameters. Quantifications show the mean value per mouse. Nested one-way ANOVA with Tukey’s multiple comparisons (2 cells per hemisphere per mice). *p<0.05, **p<0.01, ***p<0.001. Data are presented as mean values ^+^/- SEM.

### The ABA model alters the central expression of glial and inflammatory markers in a region-specific manner

We next studied the mRNA expression levels encoding glial and inflammatory markers in the hippocampus and amygdala. Globally, a group effect was revealed in both acute and chronic models (Additional file 3 and 4) that was tissue- and time-dependent. The mRNA expression of astrocytic-associated transcripts (*Gfap, S100β, Slc1a2, Gja1*) was decreased in the hippocampus of acute and chronically restricted mice compared to CT (Fig.5b and 5e). In the amygdala, no differences were observed in the acute model (Fig.5c). By contrast, in the chronic model, a downregulation of *Gfap* mRNA was observed in LFA and ABA, as well as an upregulation of *S100β* and *Slc1a2* in ABA mice (Fig.5f). Regarding the microglial markers, the mRNA expression of *Aif1, Itgam, Tmem119* and *Trem2* was decreased in the hippocampus in both acute and chronic restricted mice (Fig.5b and 5e). *Cx3cr1* mRNA level was reduced in restricted mice in the acute model while it remained unchanged in the chronic models. By contrast, a significant decrease in *P2ry12* mRNA expression was only observed in the chronic ABA group (p<0.01 vs CT, Fig.5e). Regarding the amygdala, a similar profile was observed in the acute restricted mice (Fig.5c), whereas microglial response showed different patterns compared to the hippocampus in the chronic model. Although *Itgam* and *Trem2* mRNA levels were reduced in the chronic restricted mice as observed in the hippocampus, *Aif1, P2ry12 and Tmem119* mRNA expression was not changed in the amygdala in the chronic ABA group. By contrast, the mRNA level for *Cd68* was increased in chronic ABA mice (p<0.01 vs CT and p<0.05 vs LFA, Fig.5f).

About the mRNA expression of pro-inflammatory markers in the hippocampus, we observed marked differences between acute and chronic models. Indeed, while acute ABA mice only exhibited a decrease in the anti-inflammatory marker, *Tgfb1* (Fig.5b), chronic ABA mice showed a decrease in *Il1b, Nos2* and *Tlr4* mRNA expression (Fig.5e). In addition, the expression of the neurotrophic marker *Bdnf* was increased in the hippocampus in chronic ABA mice (Fig. 5e), while only a trend for an increase was observed in acute ABA mice (Fig.5b). Similarly, in the amygdala, no significant changes were highlighted in acute mice (Fig.5c). By contrast, in the chronic ABA group, the *Nos2* and *Tgfb1* mRNA expression was reduced compared to control mice, whereas the mRNA expression of *Tnf* was increased (Fig.5f).

**Fig. 5:**
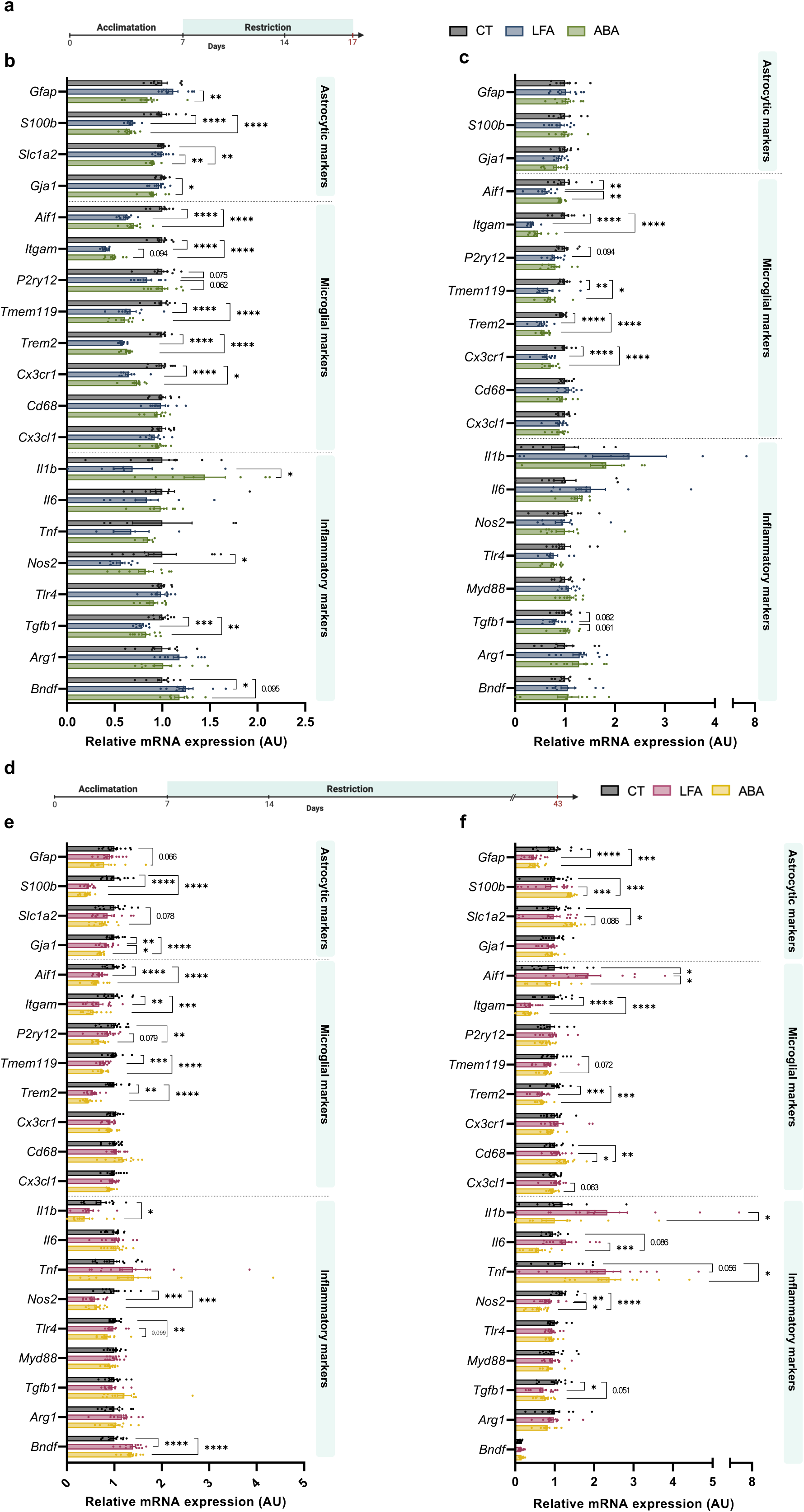
**Acute and chronic ABA models alter glial and inflammatory mRNA markers in a region-specific manner.** Female C57BL/6J mice underwent a quantitative restriction of 60% of the individual reference value of food intake for 1 week, to reach a 25% body weight loss, then an adjusted 50% restriction to maintained it for 3 days (acute; n=27) or until 43 days (chronic; n=36). Mice had free access to a monitored running wheel (ABA; n=9 for acute, n=12 for chronic) or not (limited-food access (LFA); n=9 for acute, n=12 for chronic). Control mice were fed ad libitum and had no access to a running wheel (CT; n=9 for acute, n=12 for chronic). **(a)** Experimental design of acute model. **(b)** Relative mRNA levels of astrocytic, microglial and inflammatory markers in the hippocampus and **(c)** amygdala, evaluated by RT-qPCR. **(d)** Experimental design of chronic model. **(e)** Relative mRNA levels of astrocytic, microglial and inflammatory markers in the hippocampus and **(f)** amygdala, evaluated by RT-qPCR Data are normalized on *Eef2* and *Rps18* reference gene expression and presented as a ratio of CT group. Ordinary one-way ANOVA with Tukey’s multiple comparisons or Kruskal-Wallis with Dunn’s post-test according to normality. *p<0.05, **p<0.01, ***p<0.001. Data are presented as mean values ^+^/- SEM.

**Fig. 6:**
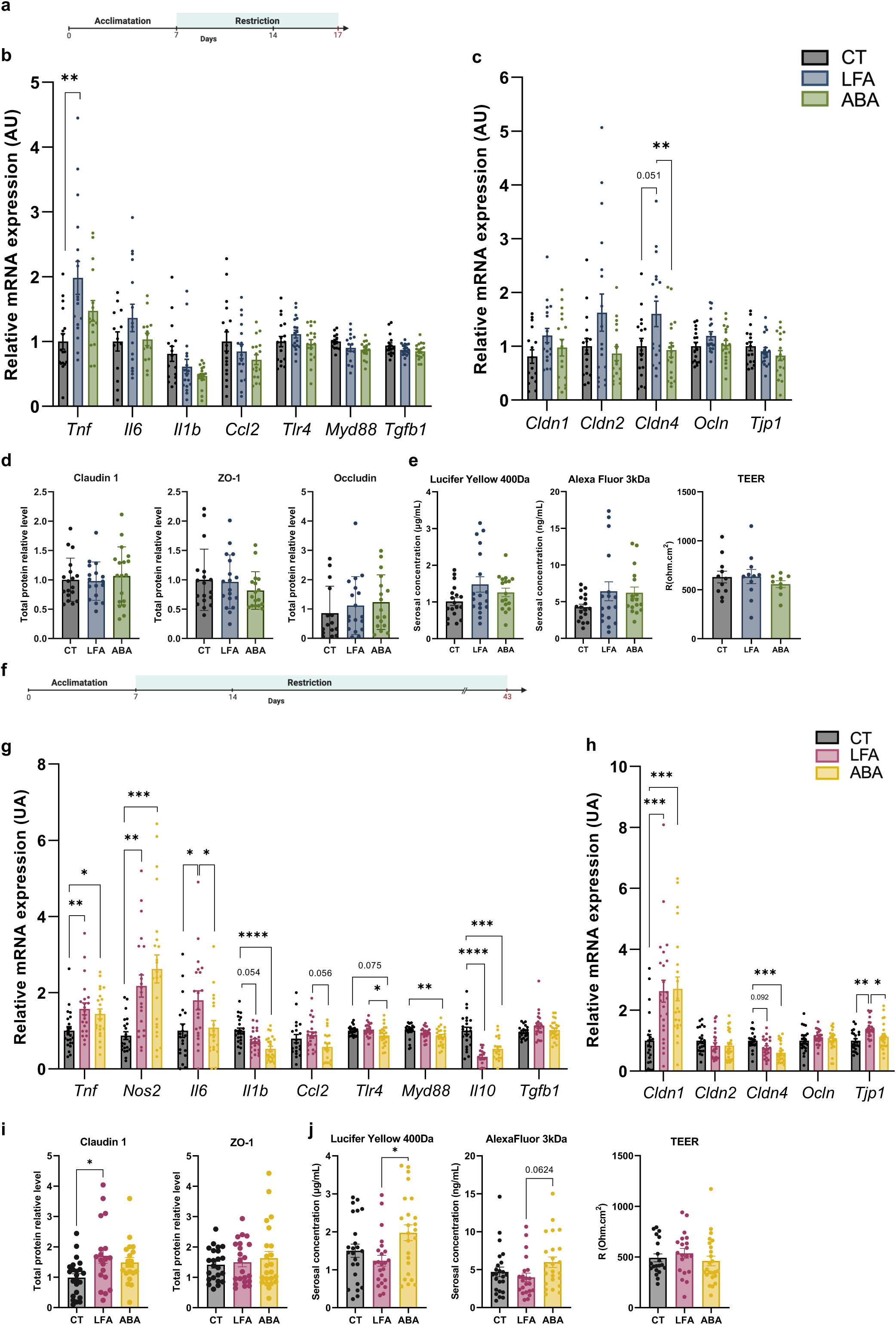
**Chronic, but not acute, ABA model induces a modulation of colonic structure, function, and inflammation.** Female C57BL/6J mice underwent a quantitative restriction of 60% of the individual reference value of food intake for 1 week, to reach a 25% body weight loss, then an adjusted 50% restriction to maintained it for 3 days (acute; n=54) or 4 weeks (chronic; n=72). Mice had free access to a monitored running wheel (ABA; n=18 for acute, n=24 for chronic) or not (limited-food access (LFA); n=18 for acute, n=24 for chronic). Control mice were fed ad libitum and had no access to a running wheel (CT; n=18 for acute, n=24 for chronic). **(a)** Experimental design of acute model. **(b)** Relative mRNA levels of colic inflammatory and **(c)** tight junction markers, evaluated by RT-qPCR. Data are normalized on *Eef2* and *Rps18* reference gene expression and presented as a ratio of CT group. **(d)** Relative protein levels of colic tight junction, normalized on total proteins and presented as a ratio of CT group, evaluated by western blot. **(e)** Serosal concentrations of Lucifer yellow and alexafluor, and trans-epithelial electric resistance (TEER) measured on a colonic fragment using Ussing chambers. **(f)** Experimental design of chronic model. **(g)** Relative mRNA levels of colic inflammatory and **(h)** tight junction markers, evaluated by RT-qPCR. Data are normalized on *Eef2* and *Rps18* reference gene expression and presented as a ratio of CT group. **(i)** Relative protein levels of colic tight junction, normalized on total proteins and presented as a ratio of CT group, evaluated by western blot. **(j)** Serosal concentrations of Lucifer yellow and alexafluor, and trans-epithelial electric resistance (TEER) measured on a colonic fragment using Ussing chambers. Ordinary one-way ANOVA with Tukey’s multiple comparisons or Kruskal-Wallis with Dunn’s post-test according to normality. *p<0.05, **p<0.01, ***p<0.001. Data are presented as mean values ^+^/- SEM.

**Fig. 7:**
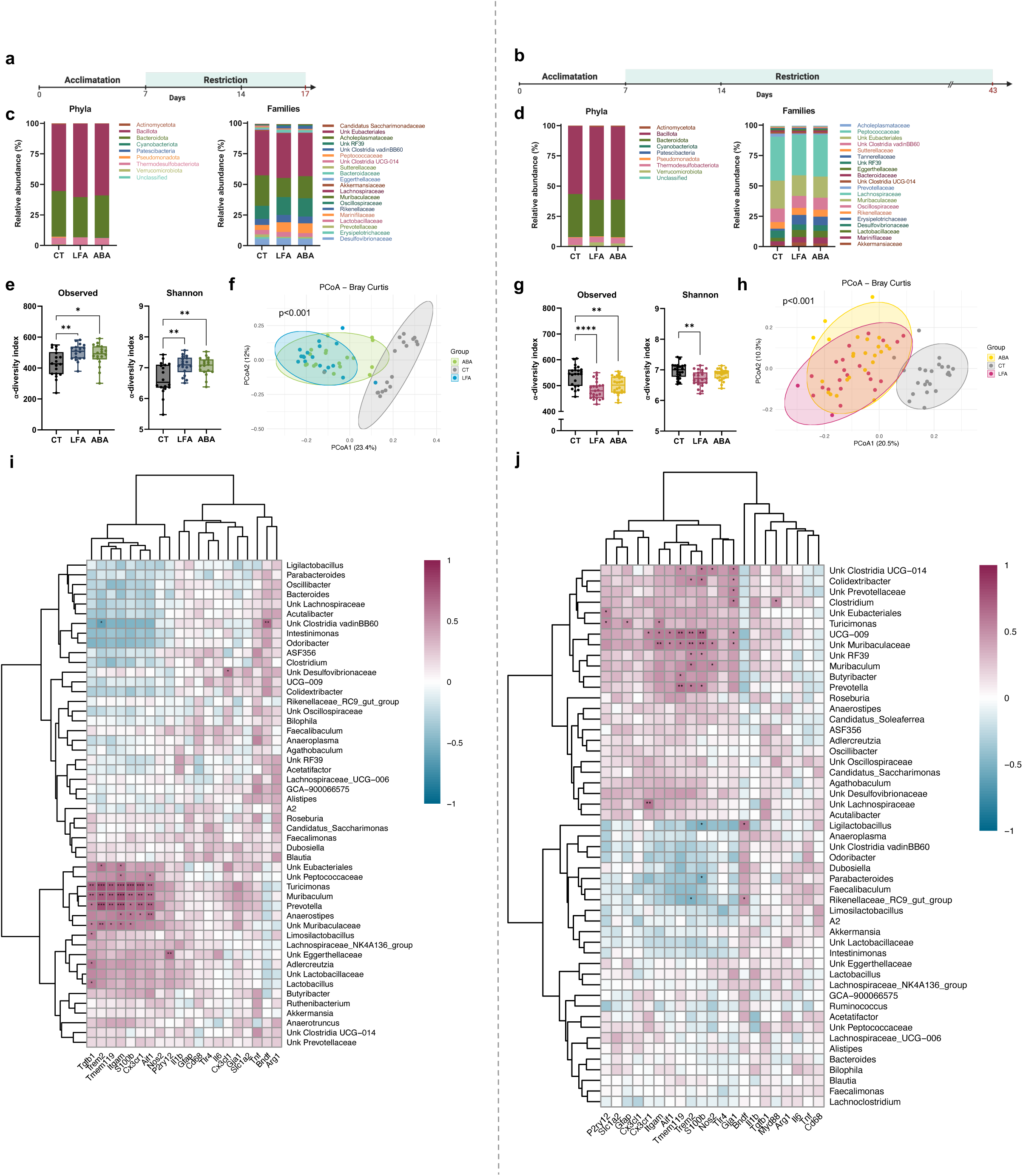
**Gut microbiota composition is impacted by food restriction in both acute and chronic ABA models.** Female C57BL/6J mice underwent a quantitative restriction of 60% of the individual reference value of food intake for 1 week, to reach a 25% body weight loss, then an adjusted 50% restriction to maintained it for 3 days (acute; n=54) or until 43 days (chronic; n=72). Mice had free access to a monitored running wheel (ABA; n=18 for acute, n=24 for chronic) or not (limited-food access (LFA); n=18 for acute, n=24 for chronic). Control mice were fed ad libitum and had no access to a running wheel (CT; n=18 for acute, n=24 for chronic). **(a)** Experimental design of acute model. **(b)** Experimental design of chronic model. **(c)** Relative abundance of bacteria phyla and the 20 most abundant families of acute and **(d)** chronic model. **(e)** lZI-diversity indexes determined by Observed and Shannon index in acute and **(g)** chronic model. Ordinary one-way ANOVA with Tukey’s multiple comparisons. Data are presented as mean values ^+^/- SEM. **(f)** Principal coordinates analysis (PCoA) of Bray-Curtis β-diversity index of acute and **(h)** chronic model. **(i)** Heatmap representation of correlations between relative mRNA levels of astrocytic, microglial and inflammatory markers in the hippocampus, and relative abundance of cecal bacteria at the genus level, in acute and **(j)** chronic model Unclassified genera are referred to as « Unk », followed by the last known taxa. The genera represented are those with an abundance greater than 0.1% across all samples and a prevalence greater than 20% in at least one group. Correlations were computed using Spearman test with Benjamini-Hochberg correction for multiple testing. Clustering was built on Euclidean distance. Bray-Curtis indexes were compared with PERMANOVA test. Cecal microbiota composition was investigated by 16S rRNA AVITI sequencing. Data are expressed as relative abundance compared to the total number of detected amplicon sequence variant (ASV). *p<0.05, **p<0.01, ***p<0.001.

**Fig. 8:**
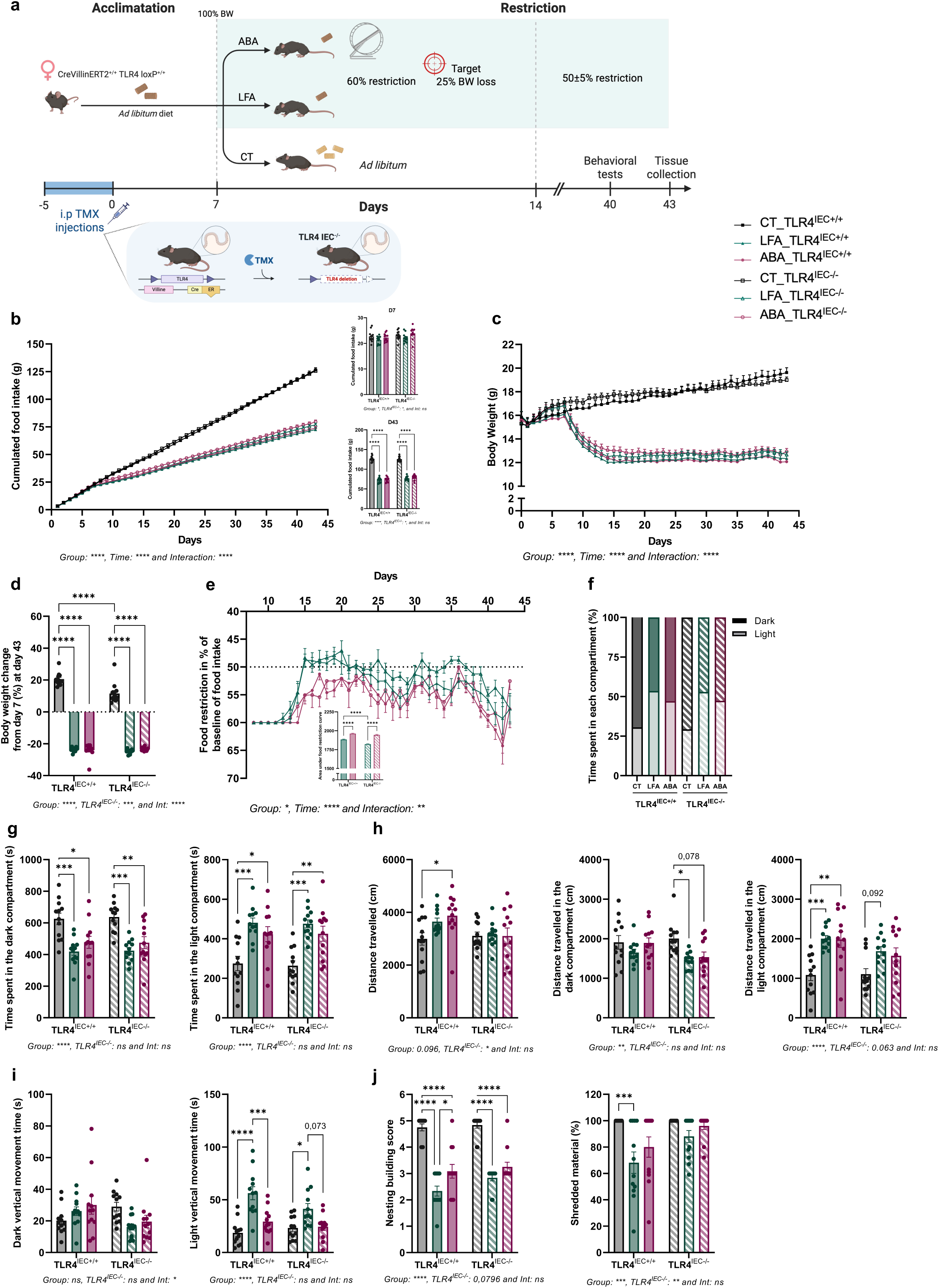
**ABA model response is partially impacted by TLR4 invalidation in IEC.** Female C57BL/6J mice underwent a quantitative restriction of 60% of the individual reference value of food intake for 1 week, to reach a 25% body weight loss, then an adjusted 50% restriction to maintained it until 43 days (n=72). Mice had free access to a monitored running wheel (ABA; n=24) or not (limited-food access (LFA); n=24). Control mice were fed ad libitum and had no access to a running wheel (CT; n=24). In each experimental group, mice are divided into invalidated (TLR4^IEC-^/^-^; n=12 per group) or not (TLR4^IEC+/+^; n=12 per group) for the TLR4 in IEC by intra-peritoneal injections of tamoxifen (TMX) before the protocol. **(a)** Experimental design. At day 40, mice were subjected to nesting and light-dark box tests **(b)** Cumulated food intake (in g) during the protocol, at D7 and D43. **(c)** Body weight (in g) throughout the protocol. **(d)** Final body weight change from day 7 (in %). **(e)** Adjustments of food restriction in percentage of reference value of food intake, and area under the curve. **(f)** Distribution of time spent in each compartment (in %). **(g)** Time (in s) spent in each compartment per group. **(h)** Distance (cm) travelled in the total device and in each compartment. **(i)** Dark and light vertical movement time (in s). **(j)** Nesting building scores and quantity of shredded material (in %) in nesting test. Ordinary two way ANOVA with Tukey’s multiple comparisons test for bar graphs, repeated measures two way ANOVA with Tukey’s multiple comparisons test for line graphs. *p<0.05, **p<0.01, ***p<0.001, ****p<0.0001. Data are presented as mean values ^+^/- SEM.

**Fig. 9:**
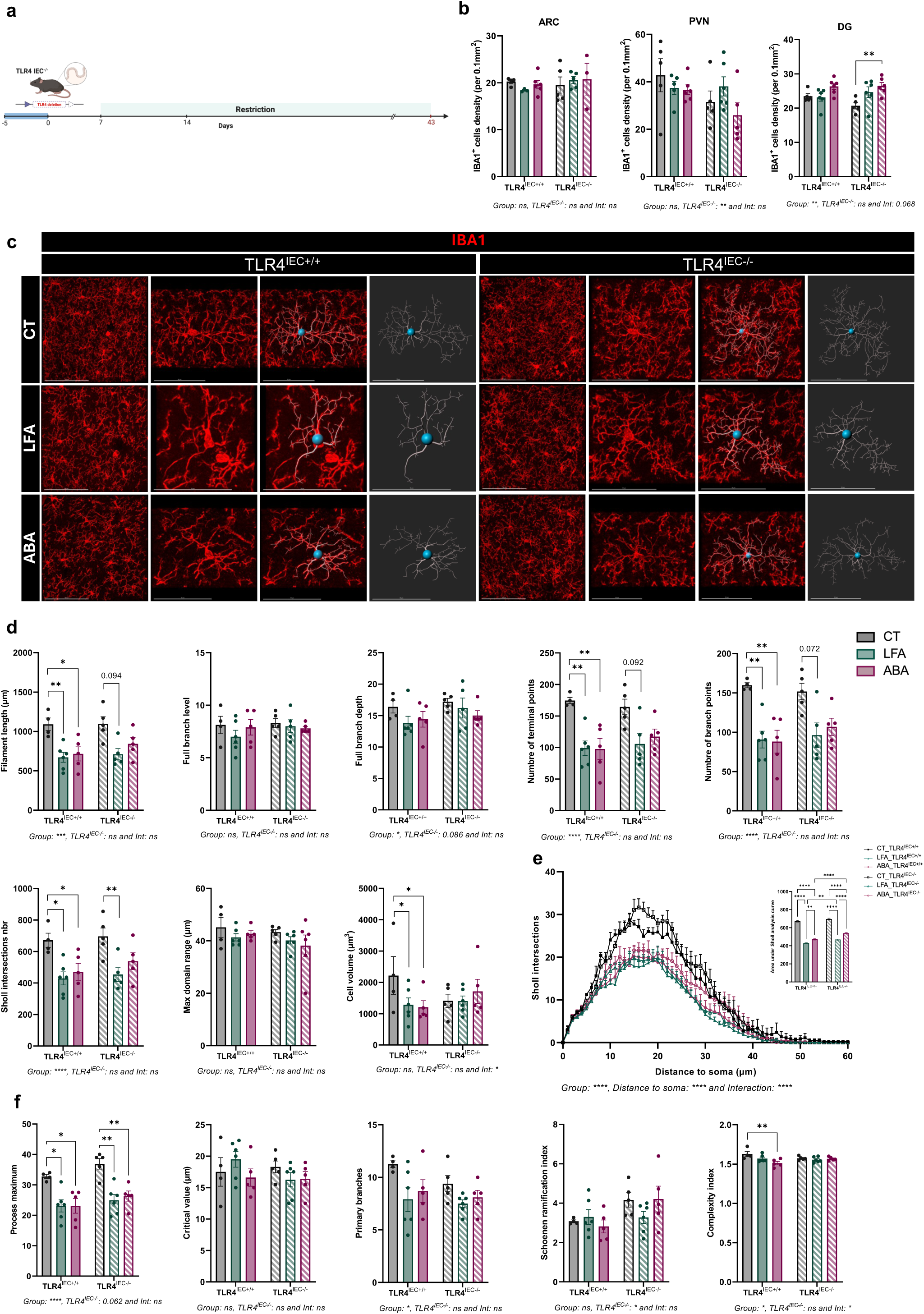
**TLR4IEC-/- impacts slightly microglial density and morphology in response to ABA model.** Female C57BL/6J mice underwent a quantitative restriction of 60% of the individual reference value of food intake for 1 week, to reach a 25% body weight loss, then an adjusted 50% restriction to maintained it until 43 days (n=36). Mice had free access to a monitored running wheel (ABA; n=12) or not (limited-food access (LFA); n=12). Control mice were fed ad libitum and had no access to a running wheel (CT; n=12). In each experimental group, mice are divided into invalidated (TLR4^IEC-^/^-^; n=6 per group) or not (TLR4^IEC+/+^; n=6 per group) for the TLR4 in IEC by intra-peritoneal injections of tamoxifen (TMX) before the protocol. **(a)** Experimental design. **(b)** IBA1^+^ cells mean number per mouse in the arcuate nucleus (ARC), paraventricular nucleus (PVN) and dentate gyrus (DG). **(c)** Confocal acquisitions of IBA1^+^ cells in the DG (scale bar: 100 μm), and zoom of a cell with morphometric reconstitution (scale bar: 50 μm). **(d)** Quantification of general morphometric parameters calculated by IMARIS software. **(e)** Number of microglial intersections over the distance to soma (Sholl analysis) and area under curve (AUC). **(f)** Quantification of Sholl parameters. Quantifications show the mean value per mouse. Nested two way ANOVA with Tukey’s multiple comparisons test for density analysis (4 to 8 replicas per mice), ordinary two way ANOVA with Tukey’s multiple comparisons test for morphometry analysis (2 cells per mice). *p<0.05, **p<0.01, ***p<0.001, ****p<0.0001. Data are presented as mean values ^+^/- SEM.

**Fig. 10:**
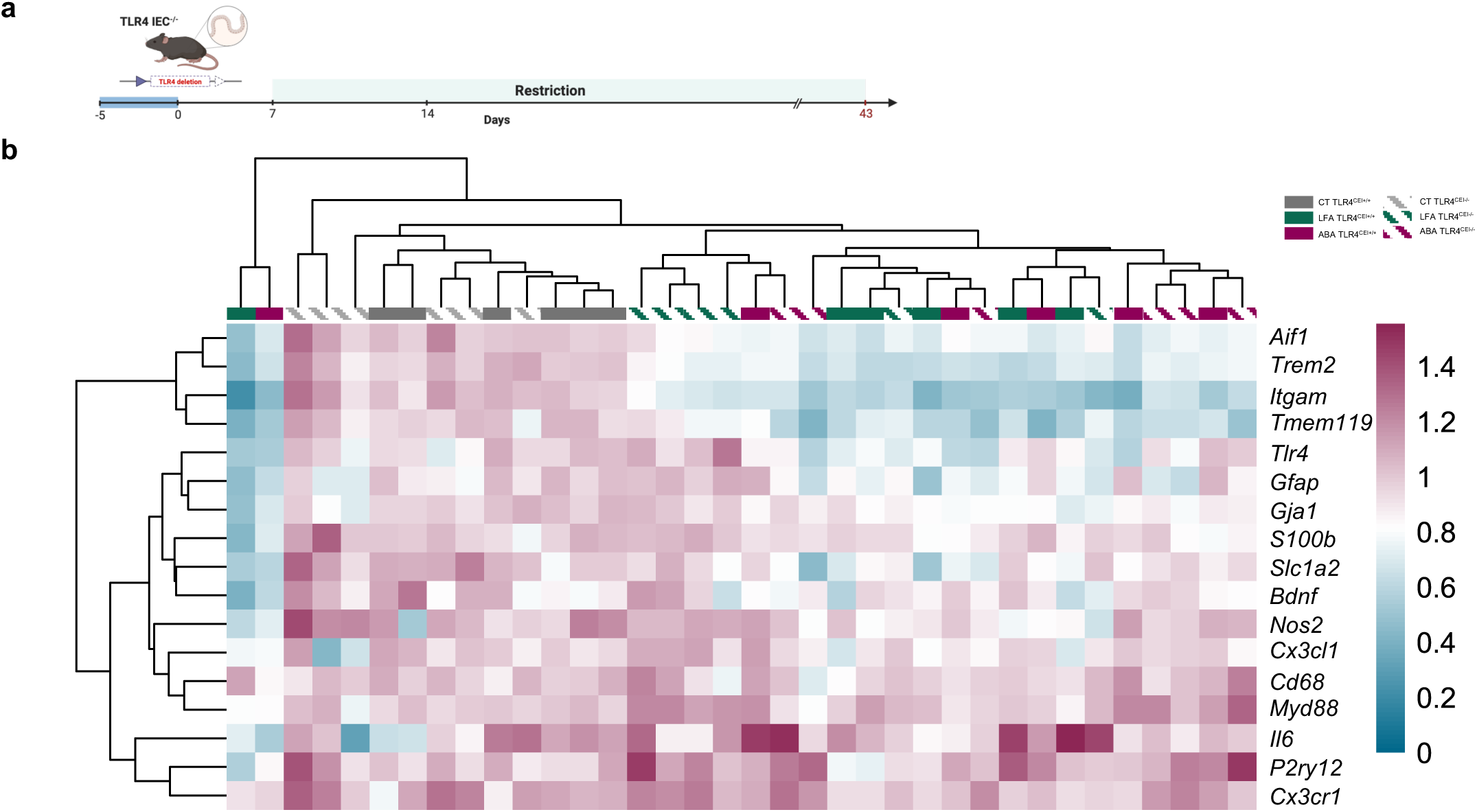
**TLR4IEC-/- impacts glial and inflammatory markers in the amygdala in response to ABA model.** Female C57BL/6J mice underwent a quantitative restriction of 60% of the individual reference value of food intake for 1 week, to reach a 25% body weight loss, then an adjusted 50% restriction to maintained it until 43 days (n=36). Mice had free access to a monitored running wheel (ABA; n=24) or not (limited-food access (LFA); n=24). Control mice were fed ad libitum and had no access to a running wheel (CT; n=24). In each experimental group, mice are divided into invalidated (TLR4^IEC-^/^-^; n=12 per group) or not (TLR4^IEC+/+^; n=12 per group) for the TLR4 in IEC by intra-peritoneal injections of tamoxifen (TMX) before the protocol. **(a)** Experimental design. **(b)** Heatmap representation of relative mRNA levels of astrocytic, microglial and inflammatory markers in the amygdala, evaluated by RT-qPCR (n=6 per group). Clustering was built on Euclidean distance. Data are normalized on *Eef2* and *Rps18* reference gene expression and presented as a ratio of CT TLR4^IEC+/+^ group. *p<0.05, **p<0.01, ***p<0.001. Data are presented as mean values ^+^/- SEM.

**Fig. 11:**
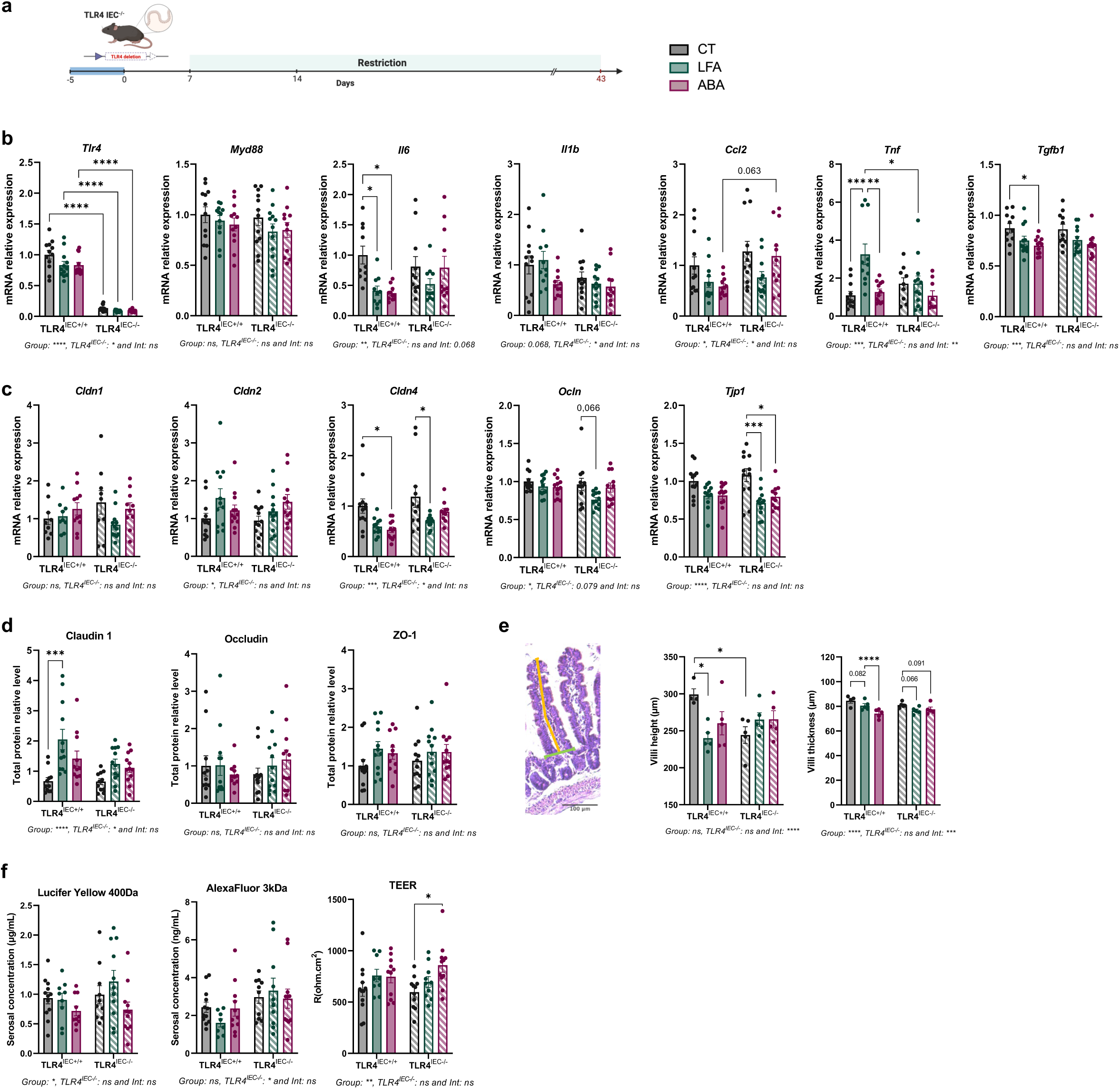
**TLR4IEC-/- induces a modulation of gut structure, function, and inflammation in response to ABA model.** Female C57BL/6J mice underwent a quantitative restriction of 60% of the individual reference value of food intake for 1 week, to reach a 25% body weight loss, then an adjusted 50% restriction to maintained it until 43 days (n=36). Mice had free access to a monitored running wheel (ABA; n=24) or not (limited-food access (LFA); n=24). Control mice were fed ad libitum and had no access to a running wheel (CT; n=24). In each experimental group, mice are divided into invalidated (TLR4^IEC-^/^-^; n=12 per group) or not (TLR4^IEC+/+^; n=12 per group) for the TLR4 in IEC by intra-peritoneal injections of tamoxifen (TMX) before the protocol. **(a)** Experimental design. **(b)** Relative mRNA levels of colic inflammatory and **(c)** tight junction markers, evaluated by RT-qPCR. Data are normalized on *Eef2* and *Rps18* reference gene expression and presented as a ratio of CT TLR4^IEC+/+^ group. **(d)** Relative protein levels of colic tight junction, normalized on total proteins and presented as a ratio of CT TLR4^IEC+/+^ group, evaluated by western blot. **(e)** Histological measures of jejunal villi height (in orange) and thickness (in green) (in μm). **(f)** Seric concentrations of Lucifer yellow and alexafluor, and trans-epithelial electric resistance (TEER) measured on a colonic fragment using Ussing chambers. Ordinary two-way ANOVA with Tukey’s multiple comparisons for **(b)** to **(d)** and *(f)*, and nested two-way ANOVA with Tukey’s multiple comparisons, on the mean of each region of interest (10 to 12 per mouse, n=5) for **(e)** . *p<0.05, **p<0.01, ***p<0.001. Data are presented as mean values ^+^/- SEM.

Interestingly, we also observed that LFA and ABA showed different responses suggesting a role of the physical activity. Particularly, in the amygdala, chronic LFA mice showed a higher mRNA expression for *Il1b* and *Il6* compared to ABA mice (Fig.5f).

### The chronic, but not acute, ABA model alters the colonic inflammation and gut barrier function

Considering the established interconnection between the brain and the gastrointestinal system, we then wanted to gain insights into gut inflammation by evaluating the expression of inflammatory markers and tight junction proteins in the colonic mucosa, and by assessing colonic gut barrier function.

In the acute model, none of the evaluated inflammatory markers were modified in ABA mice compared to control mice (Fig.6b). By contrast, the chronic model led to more numerous changes in inflammatory targets. *Tnf* and *Nos2* mRNA levels were increased in chronic ABA mice compared to CT, while *Il1b, Myd88* and *Il10* mRNA levels were reduced (Fig.6g). LFA mice showed specific response with an increase of *Tnf* and *Il6* mRNA at the acute and chronic phases, respectively.

Gut barrier function has been evaluated by qPCR and Western-blotting directed against tight junction proteins and by measuring flux levels of fluorescent molecules and electric resistance on colonic samples placed in Ussing chambers. In the acute model, ABA mice did not show significant changes of colonic barrier function since neither tight junction protein levels nor colonic para-cellular permeability were altered (Fig.6c-e). By contrast, in the chronic ABA model, *Cldn1* transcript was upregulated in both restricted mice, whereas mRNA transcripts of *Cldn4* were decreased in ABA compared to CT mice (Fig.6h). Tight junction protein levels remained unaffected in ABA mice (Fig.6i). Concerning colonic permeability, higher serosal concentrations were measured in chronic ABA mice compared to LFA, with both Lucifer yellow and AlexaFluor dyes (p<0.05 and p=0.062, respectively, Fig.6j), but electric resistance remained unchanged.

### Gut microbiota composition is affected by food restriction in both acute and chronic ABA models

To investigate how acute and chronic ABA models affect cecal microbiota composition, 16S rRNA sequencing was performed on DNAs extracted from mouse cecal contents. Overall microbiota composition differed between CT and restricted mice, in both acute and chronic models (Fig.7c and d). Acute restricted mice showed a higher lZI-diversity, in terms of richness and evenness (Fig.7e), although in chronic restricted mice we found less ASVs, and a decrease in Shannon index (Fig.7g). Bray-Curtis analysis of β-diversity highlighted distinct bacterial communities between restricted and CT mice whereas ABA and LFA microbiota composition remain very similar in both acute (p<0.001, Fig.7f) and chronic models (p<0.001, Fig.7h).

At the phylum level, restricted mice from the acute model displayed a decrease in *Actinomycetota* and *Pseudomonadota* relative abundances (Additional file 5c). In the chronic model, we identified an increase in *Bacillota* and a decrease in *Bacteroidota*, *Thermodesulfobacteriota* and *Cyanobacteriota* relative abundances between ABA and CT mice (Additional file 5d).

The main changes occurring at the family level are summarized in Additional file 6d. Briefly, an increase in *Marinifilaceae* and *Clostridia vadinBB60* and a decrease in *Muribaculaceae* and *Prevotellaceae* abundances were found in both acute and chronic models. Some specificities were observed in each model, especially an increase in *Erysipelotrichaceae* and *Lactobacillaceae* and a decrease in unknown *RF39* and *Desulfovibrionaceae* in chronic LFA and ABA mice. In addition, ABA chronic mice displayed a lower abundance of unknown *Clostridia UCG-014* and *Eubacteriales* (Additional file 5e and f).

### Gut microbiota taxa are correlated with glial markers and morphometry

We next explored the correlations between the previous microbiota changes, at the genus level, and the expression of glial and inflammatory mRNA markers in the brain, by Spearman’s test. In the hippocampus of the acute model (Fig.7i), a cluster composed of *Turicimonas*, *Muribaculum*, *Prevotella*, *Anaerostipes*, unknown *Muribaculaceae, Eubacteriales* and *Peptococcaceae* was positively correlated with some glial markers as *Aif1*, *S100b*, *Trem2*, *Tmem119*, *Cx3cr1* and *Itgam*. A few similarities were retrieved in the amygdala, especially the correlation between *Turicimonas*, *Muribaculum* and *Prevotella,* and *Cx3cr1* and *Itgam* (Additional file 5g).

In the hippocampus of the chronic model (Fig.7j), *Turicimonas*, *Muribaculum*, *Prevotella*, *Anaerostipes*, unknown *Muribaculaceae, Eubacteriales* and *Peptococcaceae* were once again positively correlated with glial markers. Some specific features of the chronic model have emerged as a cluster of *Butyribacter*, *UCG-009*, *Clostridium*, *Colidextribacter* and unknown *RF39*, *Prevotellaceae* and *Clostridia UCG-014* positively correlated with glial markers. In addition, *Bdnf* was positively associated with *Rikenellaceae RC9 gut group* and *Ligilactobacillus*. These genera were also negatively correlated with *Trem2* and *S100b* expression, respectively. Regarding the amygdala markers of the chronic model, *Colidextribacter*, *UCG-009*, and unknown *Muribaculaceae* were once again highlighted as positively correlated with glial markers (Additional file 5h).

We then investigate correlations between the level of gut microbiota taxa and morphometric parameters of glial cells in the dentate gyrus of chronic mice. Interestingly, in microglia, several previously identified genera were highlighted significant again (Additional file 6b). *Turicimonas*, *Muribaculum*, *Prevotella*, *Butyribacter*, unknown *Muribaculaceae* and *Clostridia UCG-014* were positively correlated with markers as the number of Sholl intersections, maximal domain range, critical value, process maximum, branch level, primary branches, and Sholl AUC. At the opposite, a cluster composed of *Ligilactobacillus*, *Rikenellaceae RC9, Limosilactobacillus,* and *Faecalibaculum* were negatively associated with the number of Sholl intersections and AUC, filament length and number of terminal and branch points. Interestingly, astrocytic morphometrics were not correlated with any cecal microbiota genera (Additional file 6c).

### TLR4 invalidation in IEC affects partially the ABA model response

Given that our results highlighted that the microbiota changes described in our models correlate with transcriptional and morphometric markers of glial cells, we wanted to explore whether altered microbiota-host crosstalk could modify the glial response to the ABA model. We thus reproduced our chronic ABA model in mice invalidated for TLR4 specifically in intestinal epithelial cells.

During the habituation phase, the cumulated food intake was slightly affected by groups and genotypes (D7, p(Group)<0.05, p(TLR4^IEC-/-^)<0.05, no post-test significances, Fig.8b), while body weight was slightly affected by TLR4 invalidation (p(TLR4^IEC-/-^)<0.01, Fig.8c and S6b) but not by experimental groups (p(Group)=ns, Additional file 7b). During the restriction phase, cumulated food intake was lower in restricted mice (p<0.0001) with only a weak effect of TLR4 invalidation in IEC (p(TLR4^IEC-/-^)<0.05, no post-test significances). After the restriction, all restricted mice reached the targeted 25% body weight loss, without differences between them (ns, Fig.8d). However, ABA mice needed a more severe food restriction than LFA to maintain the target body weight (p<0.0001, Fig.8e). TLR4 invalidation did not impact this parameter in ABA mice but LFA TLR4^IEC-/-^ group exhibited a less pronounced required-restriction than LFA TLR4^IEC+/+^ group (p<0.0001).

### TLR4^IEC^ invalidation induces a weak cell-specific impact on glial morphology in DG

Glial cell density was not affected by TLR4^IEC^ invalidation in the ARC and in the DG (Fig.9b and S7b). In the PVN, TLR4 deletion effects were observed in both IBA1^+^ and GFAP^+^ cells densities (p<0.01 and p<0.05, respectively) without post-test significances (Fig.9b and Additional file 8b).

We wanted to decipher whether TLR4^IEC^ invalidation could affect the previously described glial morphometry response in the DG. Morphological parameters and Sholl analysis associated values were globally not altered by TLR4 invalidation in IEC in the DG. However, in microglia, the decrease of parameters observed in LFA and ABA wild-type mice were not always retrieved in TLR4^IEC-/-^ mice (Fig.9d). The area under Sholl analysis curves were increased by TLR4^IEC^ invalidation in LFA and ABA mice (p<0.01 and p<0.0001, respectively, Fig.9e). Process maximum and Schoenen ramification index showed global TLR4 invalidation effects (p=0.062 and p<0.05, respectively, Fig.9f) without post-test significances. In astrocytes, the increase in the area under Sholl analysis curves in LFA and ABA wild-type mice was not observed in TLR4^IEC-/-^ mice that could be explained by the effect of TLR4^IEC^ invalidation in CT groups (p<0.0001, Additional file 8e). Similarly, the increase of the Schoenen ramification index in LFA wild-type mice disappeared in TLR4^IEC-/-^ mice (Additional file 8f).

### TLR4 invalidation in IEC impacts the central inflammatory response in the amygdala

In the hippocampus, TLR4^IEC^ invalidation did not induce strong effects on glial and inflammatory markers expression (Additional file 9). However, in the amygdala, additionally to a global group effect (Fig10b), specific changes were observed. TLR4^IEC^ invalidation was associated with significant changes in *Slc1a2, Itgam, P2ry12*, *Trem2 Aif1* and *Cx3cr1* mRNA levels (Additional file 10b). Particularly, TLR4^IEC^ invalidation induced an increase of *Aif1* expression in LFA mice (p<0.05) and *Cx3cr1* in CT mice (p<0.05). In addition, *Gfap* and *Tmem119* were increased after TLR4^IEC^ invalidation in LFA mice (p<0.05) but not in ABA mice.

### Colonic inflammation and gut barrier function are slightly modified by TLR4 invalidation in IEC

mRNA transcripts encoding inflammatory markers were evaluated in the colon to assess the involvement of TLR4^IEC^ in inflammatory responses observed in the chronic ABA model (Fig.11b). We first confirmed the invalidation of *Tlr4* in IEC in the concerned groups. Then, the increased expression of *Tnf* in LFA mice was not observed in TLR4^IEC-/-^ subgroup with a downregulation in invalidated LFA compared to wild-type LFA (p<0.05, p(Interaction)<0.01). A TLR4^IEC-/-^ effect was also found in *Il1b* and *Ccl2* mRNA levels (p(TLR4^IEC-/-^)<0.05) with a trend for an upregulation of *Ccl2* in ABA TLR4^IEC-/-^ compared to wild-type ABA (p=0.063).

Interestingly, the decrease in *Il6* and *Tgfb1* mRNA levels observed in wild-type ABA mice compared to wild-type control mice disappeared in TLR4^IEC-/-^ mice.

We assessed mRNA and protein levels of tight junction proteins in the colon. TLR4 invalidation in IEC globally affected *Cldn4* mRNA expression (p(TLR4^IEC-/-^)<0.05, Fig.11c) without post-test significances. Additionally, *Tjp1* mRNA expression was significantly downregulated in restricted TLR4^IEC-/-^ mice (Fig.11c) but not in wild-type mice. Tight junction protein levels revealed an invalidation effect in claudin-1 expression (p(TLR4^IEC-/-^)<0.05, Fig.11d). The increase in claudin-1 observed in wild-type LFA and ABA mice were not retrieved in TLR4^IEC-/-^ subgroups (Fig.11d). Concerning the colonic *ex vivo* permeability assessed using Ussing chambers, we observed that serosal concentration of AlexaFluor showed a global TLR4^IEC^ invalidation effect (p<0.05), while serosal concentration of Lucifer yellow and electric resistance were not affected (Fig.11f).

### Food restriction impacts jejunal villi morphology

We also investigated jejunal villi morphology by carried out H&E staining. Villi height displayed only post-test significances with shorter villi in LFA wild-type mice compared to CT (p<0.05, Fig.11e), and in CT TLR4^IEC-/-^ compared to CT wild-type (p<0.05). Villi were thinner in response to restriction (p(Group)<0.001), depending on genotype (p(TLR4^IEC-/-^)<0.001), with post-test significances or tendencies between restricted mice and their associated CT (p=0.082 for LFA TLR4^IEC+/+^ and p<0.0001 for ABA TLR4^IEC+/+^ vs CT TLR4^IEC+/+^, p=0.066 for LFA TLR4^IEC-/-^ and p=0.091 for ABA TLR4^IEC-/-^ vs CT TLR4^IEC-/-^).

### Restricted mice show altered exploratory and anxiety behavior, with minor effects of TLR4^IEC^ invalidation

We next performed light-dark box test to assess unconditioned anxiety response and spontaneous exploratory behavior of mice. First, regardless of genotype, CT mice spent more time in the dark compartment than in the light one (Fig.8f), which allows us to validate the correct operation of the test. Interestingly, restricted mice did not seem to dissociate the two compartments, with the same amount of time spent in both (Fig.8f), without any impact of TLR4^IEC^ invalidation. Restricted wild-type mice showed a higher total distance travelled and light distance travelled than control mice (Fig.8h). This difference was partially reduced in TLR4^IEC-/-^ mice. By contrast, TLR4^IEC-/-^ restricted mice showed a lower distance travelled in dark compartment than control mice that was not observed in wild-type mice (Fig.8h). We found an increase in the time spent in vertical movements in LFA mice specifically, in both genotypes, in the light compartment but not in the dark one (Fig.8i). We also evaluated the anxiety response by using nesting test. We only observed an effect of TLR4 invalidation in IEC for shredded material (p(TLR4^IEC-/-^)<0.01, Fig.8j) without post-test significances. The response observed in wild-type LFA mice was not retrieved in TLR4^IEC-/-^ mice.

## Discussion

The aim of this work was to evaluate the neuroinflammation and its behavioral consequences in the ABA model at the acute and chronic phases, and to evaluate the role of the microbiota-gut-brain axis in these alterations. We show that the chronic ABA model alters glial response, particularly in the hippocampus, and is associated with gut microbiota changes. However, these alterations are unaffected by TLR4 invalidation in intestinal epithelial cells, suggesting other underlying mechanisms.

We first evaluated the glial response in the main brain regions involved in eating behavior and found that the chronic ABA model induced more alterations than the acute and particularly in the DG of the hippocampus, with an increase in IBA1^+^ cells number and a decrease in GFAP^+^ cells. We thus focused on this region to perform morphometric analysis, and we observed that glial cells react to food restriction by adopting a specific phenotype, with a cell-specific response once again. Indeed, microglia exhibited a loss of complexity and a global deramification in both restricted groups, although astrocytes were impacted only in LFA mice. Our results reinforce the previous hypothesis that an energy restriction could impact astrocytes by reducing their neogenesis(41) and by inducing morphological changes(21,42). These results could also explain the loss of cerebral volume found in patients with AN(43) and related preclinical model(18). Studying astrocyte-neuron coupling would reveal the potential impact of this astrocytic alteration on neuronal activity. Microglial responses can be very heterogeneous depending on the type and intensity of restriction and the brain region studied. For example, a deramification was observed in the medial prefrontal cortex, orbitofrontal cortex(44) and hippocampus(22) in a model of dehydration-induced anorexia in rats, whereas an increase in process length was found in the cortex of chronic ABA mice(45). As microglial function should not be inferred from morphology or gene expression(46), we were unable to precise if the response we observed is beneficial or detrimental. Serum neurofilament light chain levels were increased in chronic starved mice(45) but this result is not sufficient to highlight a causal effect of microglial response in neurodegeneration. Blocking microglial morphology remodeling with minocycline or by inhibiting Cdk-1 to gain insight into the role of its response could be of interest(47). Our results provide evidence for a neuroinflammatory response in the hippocampus, as previously suggested in other models(17,22). The hippocampus is well known to be implicated in eating behavior as part of the mesolimbic system, where it regulates associative learning between the meal stimuli and post-ingestive consequences, but hippocampal neurons are also activated by internal cues, such as signals transmitted by the vagus nerve or hormones, as it expresses many food-related receptors(48). Thus, perturbations in the functioning of the hippocampus are quite relevant in the AN context and may contribute to the altered eating behavior and cognition observed in patients(43). Interestingly, the amygdala responded differently to the hippocampus, with an increase in the expression of some astrocytic and proinflammatory markers in the chronic model. The amygdala is involved in emotional processes linked to conditioned learning and food intake. In patients with AN, the volume of several amygdala nuclei were reduced(49) and its activity is increased and correlated to the feeling of disgust(50). This tissue-specific response emphasizes the complexity of the brain reaction to the model. We did not observe a strong inflammatory response in the acute model, suggesting the importance of the duration of the food restriction leading to the establishment of inflammatory responses. A more precise kinetic response could be further investigated, given that the glial response observed in obesity models seems to precede the peripheral inflammatory response(51). These data were associated to altered behavioral responses in ABA mice. Indeed, we also performed behavioral tests to assess the impact of chronic ABA model under unconditioned anxiety response and spontaneous exploratory behavior. Nesting test suggested a higher anxiety state in restricted mice. In light-dark box test, restricted mice seemed to not dissociate the two compartments and show more rearing (vertical movements) in the illuminated one, that can be interpreted as abnormal behavior or stereotypy, or also foraging behavior.

Considering the established interconnection between the brain and the gastrointestinal system and the role of the microbiota-gut-brain axis, we assessed gut barrier function and inflammation in ABA mice, and showed that chronic ABA model is also associated with a colonic inflammation and a gut barrier dysfunction, as previously shown in other models(8,9) and in patients with AN(6,10). Gut microbiota composition also appeared affected by food restriction and some genera were identified as correlated with glial markers and morphometry, especially with microglia. Gut microbiota can impact microglia development and functioning through its metabolites, especially short-chain fatty acids, acting along the gut-brain axis(52). Thus, the dysbiosis described in ABA mice may be able to influence glial cells. We thus wanted to explore whether altered microbiota-host crosstalk could modify the glial response to the ABA model, by reproducing the chronic ABA model in mice specifically and conditionally invalidated for the TLR4 in intestinal epithelial cells. In previous studies, whole body TLR4 KO was associated to a high rate of mortality in the ABA model with timely-restricted food access(8), while behavioral alterations were observed after TLR4^IEC^ invalidation(14). However, in these latter studies, central response was not deeply assessed. In the present study, we globally did not observe a strong effect of TLR4^IEC^ invalidation on the studied parameters in ABA mice, even if TLR4^IEC^ invalidation induced some effects in the amygdala. Similarly, anxiety and exploratory behaviors were not altered by TLR4^IEC^ invalidation. Thus, the neuroinflammatory process observed in ABA mice are not explained by the activity of TLR4^IEC^, but it would be interesting to further evaluate the involvement of TLR4 in other cell types such as microglia. Indeed, TLR4 is expressed most abundantly by microglial cells among brain cells. Interestingly, microglia-specific TLR4 KO reversed neuroinflammatory response to a high-fat diet in a region and sex-specific manner(53).

We carried out the ABA model with quantitative food restriction as previously described(24,25) to induce a 25% body weight loss, which was maintained in an acute or chronic protocol. Maintaining the target body weight throughout the chronic protocol required an increase in food restriction, suggesting an increasing resistance to body weight loss during the protocol. In the ABA model with timely-restricted food access, a study highlighted a body weight gain in restricted mice at the end of the protocol and hypothesized about a behavioral adaptation(8). As food intake was controlled by the experimenter, our results suggest rather a metabolic adaptation, that could be deciphered in further experiments using metabolic and calorimetric cages. Finally, we also observed a small effect of physical activity, reflected by differences between LFA and ABA mice. We notably found that ABA mice seem more resistant to food restriction than LFA, as they need a more severe restriction to achieve the targeted body weight loss. In addition, LFA mice were more anxious, and exhibited astrocytic deramification. Thus, physical activity could be beneficial in our model, that could be explained by metabolic adaptations. For instance, a previous study reported a shift to lipid oxidation for LFA mice but not for ABA mice(25). In contrast, this effect was not related to gut microbiota since we did not observe physical activity-induced gut microbiota alterations, as previously shown(11). However, physical activity has been shown to differently affect gut microbiota in several studies according to the type of exercise(54–56), but the subjects were not submitted to an energetic challenge in these studies. In the present study, physical activity was only measured via monitoring the activity in the running wheels. To confirm our hypothesis, an actimeter device should be used to measure total activity. Finally, these results also support the use of adapted physical activity during refeeding for patients with AN(57).

## Conclusion

Our results reveal that chronic restriction is more strongly associated with gut inflammation, gut microbiota alterations and neuroinflammatory processes in the hippocampus than acute restriction. Microglial cells show a global deramification and loss of complexity that correlates with alterations in the level of specific gut microbiota taxa.

However, we show that intestinal epithelial TLR4 is not involved in these alterations. Our data offer a more integrative view of the consequences of the ABA model, and more generally of food restriction on mouse physiology. All together, our results highlight the negative impact of an extended food restriction and thus underly the importance of a faster diagnosis and medical care. The contribution of the digestive system, gut microbiota, and central nervous system has also been demonstrated, highlighting the importance of redefining AN in a more integrated way.

## List of abbreviations

ABA: activity-based anorexia
ACC: accumbens nucleus
AN: anorexia nervosa
ARC: arcuate nucleus
ASV: amplicon sequence variant
AUC: area under curve
CA3: cornu ammonis 3
CT: control
DAPI: 4’ ,6-diamidino-2-phenylindole
DG: dentate gyrus
DNA: desoxyribonucleic acid
i.p.: intra-peritoneal
IEC: intestinal epithelial cells
LFA: limited-food access
PBS: phosphate-buffered saline
PCoA: principal coordinates analysis
PVN: paraventricular nucleus
RNA: ribonucleic acid
ROI: region of interest
RTqPCR: reverse transcription quantitative polymerase chain reaction
SEM: standard error of the mean
TEER: trans-epithelial electric resistance
TLR4: toll-like receptor 4
TMX: tamoxifen

## Declarations

### Ethical approval and consent to participate

Not applicable.

### Consent for publication

**Not applicable.**

### Availability of data and materials

The datasets used and/or analysed during the current study are available from the corresponding author on reasonable request.

## Authors’ contributions

Conceptualization, LR and MC; LR, TD, CQ, LC, DR and MC, formal analysis; LR, TD, CS, CQ, CG, CBF, OM, AT, FL, JCdR, JLdR, and LL, investigation; LR and MC, original draft preparation; all authors, writing review and editing.

## Acknowledgements

We thank the PRIMACEN platform (HeRaCleS Inserm US51, CNRS UAR 2026, Université de Rouen Normandie, France) for the microscopy facilities.

---

**Additional file 1: Primers sequences used in RT-qPCR.**

**Additional file 2: Acute and chronic ABA models induce specific body weight loss and physical activity patterns.**

Female C57BL/6J mice underwent a quantitative restriction of 60% of the individual reference value of food intake for 1 week, to reach a 25% body weight loss, then an adjusted 50% restriction to maintained it for 3 days (acute; n=54) or 4 weeks (chronic; n=72). Mice had free access to a monitored running wheel (ABA; n=18 for acute, n=24 for chronic) or not (limited-food access (LFA); n=18 for acute, n=24 for chronic). Control mice were fed ad libitum and had no access to a running wheel (CT; n=18 for acute, n=24 for chronic). **(a)** Experimental design of acute model. **(b)** Experimental design of chronic model. **(c)** Body weight change from reference value of body weight (in %), in acute model. **(d)** Body weight change from reference value of body weight (in %), in chronic model. **(e)** wheel activity pattern (in km), in acute model. **(f)** wheel activity pattern (in km), in chronic model. Repeated measures two way ANOVA with Tukey’s multiple comparisons. #p<0.05, ##p<0.01, ###p<0.001, ####p<0.0001 ABA vs LFA. Data are presented as mean values ^+^/- SEM.

**Additional file 3: Acute ABA model alters glial and inflammatory mRNA markers expression in a region-specific manner.**

Female C57BL/6J mice underwent a quantitative restriction of 60% of the individual reference value of food intake for 1 week, to reach a 25% body weight loss, then an adjusted 50% restriction to maintained it for 3 days (n=27). Mice had free access to a monitored running wheel (ABA; n=9) or not (limited-food access (LFA); n=9). Control mice were fed ad libitum and had no access to a running wheel (CT; n=9). **(a)** Experimental design of acute model. **(b)** Heatmap representation of relative mRNA levels of astrocytic, microglial and inflammatory markers in the hippocampus and **(c)** amygdala, evaluated by RT-qPCR. Clustering was built on Euclidean distance. Data are normalized on *Eef2* and *Rps18* reference gene expression and presented as a ratio of CT group.

**Additional file 4: Chronic ABA model alters glial and inflammatory mRNA markers expression in a region-specific manner.**

Female C57BL/6J mice underwent a quantitative restriction of 60% of the individual reference value of food intake for 1 week, to reach a 25% body weight loss, then an adjusted 50% restriction to maintained it until 43 days (n=36). Mice had free access to a monitored running wheel (ABA; n=12) or not (limited-food access (LFA); n=12). Control mice were fed ad libitum and had no access to a running wheel (CT; n=12). **(a)** Experimental design of chronic model. **(b)** Heatmap representation of relative mRNA levels of astrocytic, microglial and inflammatory markers in the hippocampus and **(c)** amygdala, evaluated by RT-qPCR. Clustering was built on Euclidean distance. Data are normalized on *Eef2* and *Rps18* reference gene expression and presented as a ratio of CT group.

**Additional file 5: Gut microbiota composition is impacted by food restriction in both acute and chronic ABA models.**

Female C57BL/6J mice underwent a quantitative restriction of 60% of the individual reference value of food intake for 1 week, to reach a 25% body weight loss, then an adjusted 50% restriction to maintained it for 3 days (acute; n=54) or until 43 days (chronic; n=72). Mice had free access to a monitored running wheel (ABA; n=18 for acute, n=24 for chronic) or not (limited-food access (LFA); n=18 for acute, n=24 for chronic). Control mice were fed ad libitum and had no access to a running wheel (CT; n=18 for acute, n=24 for chronic). **(a)** Experimental design of acute model. **(b)** Experimental design of chronic model. **(c)** Relative abundance of bacteria phyla in acute and **(d)** chronic models. **(e)** Relative abundance of the 20 most abundant families in acute and **(f)** chronic models. Phyla and families were compared using ordinary one-way ANOVA with Tukey’s multiple comparisons or Kruskal-Wallis with Dunn’s post-test according to normality, corrected by Benjamini-Hochberg for multiple testing. The less abundant families are referred to as « others ». **(g)** Heatmap representation of correlations between relative mRNA levels of astrocytic, microglial and inflammatory markers in the amygdala, and relative abundance of cecal bacteria at the genus level, in acute and **(h)** chronic models. Unclassified genera are referred to as « Unk », followed by the last known taxa. The genera represented are those with an abundance greater than 0.1% across all samples and a prevalence greater than 20% in at least one group. Correlations were computed using Spearman test and Benjamini-Hochberg corrections were applied for multiple testing. Clustering was built on Euclidean distance. *p<0.05, **p<0.01, ***p<0.001. Cecal microbiota composition was investigated by 16S rRNA AVITI sequencing. Data are expressed as relative abundance compared to the total number of detected amplicon sequence variant (ASV).

**Additional file 6: Gut microbiota composition is correlated with glial morphometric parameters in the chronic ABA model.**

Female C57BL/6J mice underwent a quantitative restriction of 60% of the individual reference value of food intake for 1 week, to reach a 25% body weight loss, then an adjusted 50% restriction to maintained it for 3 days (acute; n=54) or until 43 days (chronic; n=72). Mice had free access to a monitored running wheel (ABA; n=18 for acute, n=24 for chronic) or not (limited-food access (LFA); n=18 for acute, n=24 for chronic). Control mice were fed ad libitum and had no access to a running wheel (CT; n=18 for acute, n=24 for chronic). **(a)** Experimental design of chronic model. **(b)** Heatmap representation of correlations between morphometric parameters of microglia and **(c)** astrocytes in the dentate gyrus, and relative abundance of cecal bacteria at the genus level. Unclassified genera are referred to as « Unk », followed by the last known taxa. The genera represented are those with an abundance greater than 0.1% across all samples and a prevalence greater than 20% in at least one group. Correlations were computed using Spearman test and Benjamini-Hochberg corrections were applied for multiple testing. Clustering was built on Euclidean distance. *p<0.05, **p<0.01, ***p<0.001. Cecal microbiota composition was investigated by 16S rRNA AVITI sequencing. Data are expressed as relative abundance compared to the total number of detected amplicon sequence variant (ASV). **(d)** Summary of identified families changes in acute and chronic models, compared to CT mice. Families in black are modified equally in both models, and families in green display specific changes in either acute or chronic.

**Additional file 7: ABA model response is partially impacted by TLR4 invalidation in IEC.** Female C57BL/6J mice underwent a quantitative restriction of 60% of the individual reference value of food intake for 1 week, to reach a 25% body weight loss, then an adjusted 50% restriction to maintained it until 43 days (n=72). Mice had free access to a monitored running wheel (ABA; n=24) or not (limited-food access (LFA); n=24). Control mice were fed ad libitum and had no access to a running wheel (CT; n=24). In each experimental group, mice are divided into invalidated (TLR4^IEC-^/^-^; n=12 per group) or not (TLR4^IEC+/+^; n=12 per group) for the TLR4 in IEC by intra-peritoneal injections of tamoxifen (TMX) before the protocol. **(a)** Experimental design. **(b)** Body weight (in g) at D7 and **(C)** D43. Ordinary two way ANOVA with Tukey’s multiple comparisons test. *p<0.05, **p<0.01, ***p<0.001, ****p<0.0001. Data are presented as mean values ^+^/- SEM.

**Additional file 8: TLR4 invalidation in IEC impacts slightly astrocytic density and morphology in response to ABA model.**

Female C57BL/6J mice underwent a quantitative restriction of 60% of the individual reference value of food intake for 1 week, to reach a 25% body weight loss, then an adjusted 50% restriction to maintained it until 43 days (n=36). Mice had free access to a monitored running wheel (ABA; n=12) or not (limited-food access (LFA); n=12). Control mice were fed ad libitum and had no access to a running wheel (CT; n=12). In each experimental group, mice are divided into invalidated (TLR4^IEC-^/^-^; n=6 per group) or not (TLR4^IEC+/+^; n=6 per group) for the TLR4 in IEC by intra-peritoneal injections of tamoxifen (TMX) before the protocol. **(a)** Experimental design. **(b)** GFAP^+^ cells mean number per mouse in the arcuate nucleus (ARC), paraventricular nucleus (PVN) and dentate gyrus (DG). **(c)** Confocal acquisitions of GFAP^+^ cells in the DG (scale bar: 100 μm), and zoom of a cell with morphometric reconstitution (scale bar: 25 μm). **(d)** Quantification of general morphometric parameters calculated by IMARIS software. **(e)** Number of astrocytic intersections over the distance to soma (Sholl analysis) and area under curve (AUC). **(f)** Quantification of Sholl parameters. Quantifications show the mean value per mouse. Nested two way ANOVA with Tukey’s multiple comparisons test for density analysis (4 to 8 replicas per mice), ordinary two way ANOVA with Tukey’s multiple comparisons test for morphometry analysis (2 cells per mice). *p<0.05, **p<0.01, ***p<0.001, ****p<0.0001. Data are presented as mean values ^+^/- SEM.

**Additional file 9: TLR4^IEC-/-^ does not impact glial and inflammatory markers in hippocampus in response to ABA model.**

Female C57BL/6J mice underwent a quantitative restriction of 60% of the individual reference value of food intake for 1 week, to reach a 25% body weight loss, then an adjusted 50% restriction to maintained it until 43 days (n=36). Mice had free access to a monitored running wheel (ABA; n=12) or not (limited-food access (LFA); n=12). Control mice were fed ad libitum and had no access to a running wheel (CT; n=12). In each experimental group, mice are divided into invalidated (TLR4^IEC-^/^-^; n=6 per group) or not (TLR4^IEC+/+^; n=6 per group) for the TLR4 in IEC by intra-peritoneal injections of tamoxifen (TMX) before the protocol. **(a)** Experimental design. **(b)** Relative mRNA levels of astrocytic, microglial and inflammatory markers in the hippocampus. Data are normalized on *Eef2* and *Rps18* reference gene expression and presented as a ratio of CT TLR4^IEC+/+^ group. Ordinary two-way ANOVA with Tukey’s multiple comparisons. *p<0.05, **p<0.01, ***p<0.001. Data are presented as mean values ^+^/- SEM. **(c)** Heatmap representation of relative mRNA levels of astrocytic, microglial and inflammatory markers in the hippocampus, evaluated by RT-qPCR. Clustering was built on Euclidean distance.

**Additional file 10: TLR4^IEC-/-^ impacts glial and inflammatory markers in amygdala in response to ABA model.**

Female C57BL/6J mice underwent a quantitative restriction of 60% of the individual reference value of food intake for 1 week, to reach a 25% body weight loss, then an adjusted 50% restriction to maintained it until 43 days (n=36). Mice had free access to a monitored running wheel (ABA; n=12) or not (limited-food access (LFA); n=12). Control mice were fed ad libitum and had no access to a running wheel (CT; n=12). In each experimental group, mice are divided into invalidated (TLR4^IEC-^/^-^; n=6 per group) or not (TLR4^IEC+/+^; n=6 per group) for the TLR4 in IEC by intra-peritoneal injections of tamoxifen (TMX) before the protocol. **(a)** Experimental design. **(b)** Relative mRNA levels of astrocytic, microglial and inflammatory markers in the amygdala, evaluated by RT-qPCR. Data are normalized on *Eef2* and *Rps18* reference gene expression and presented as a ratio of CT TLR4^IEC+/+^ group. Ordinary two-way ANOVA with Tukey’s multiple comparisons. *p<0.05, **p<0.01, ***p<0.001. Data are presented as mean values ^+^/- SEM.

