## Supplemental materials for "Chronic Activity-Based Anorexia triggers a glial response in the hippocampus independent of intestinal epithelial Toll-Like Receptor 4"

| Target gene | Sense | Oligonucleotides | Annealing temperature (°C) |
|---|---|---|---|
| ***Rps18*** | Forward | TGCGAGTACTCAACACCAACA | 60.0 |
| | Reverse | TTCCTCAACACCACATGAGC | |
| ***Eef2*** | Forward | TGTCAGTCATCGCCCATGTG | 60.0 |
| | Reverse | CATCCTTGCGAGTGTCAGTGA | |
| ***Aif1*** | Forward | ATCAACAAGCAATTCCTCGATGA | 61.0 |
| | Reverse | CAGCATTCGCTTCAAGGACATA | |
| ***Itgam*** | Forward | ATGGACGCTGATGGCAATACC | 62.0 |
| | Reverse | TCCCCATTCACGTCTCCCA | |
| ***Arg1*** | Forward | GTTCCCAGATGTACCAGGATTC | 61.0 |
| | Reverse | CGATGTCTTTGGCAGATATGC | |
| ***P2ry12*** | Forward | AAGTTCCCAAAGCCCTCTGT | 60.0 |
| | Reverse | ACGGACACTTTCCCGTATCC | |
| ***Cx3cr1*** | Forward | GTTATTTGGGCGACATTGTGGC | 61.5 |
| | Reverse | CAGACCGAACGTGAAGACGAG | |
| ***Cd68*** | Forward | TGTCTGATCTTGCTAGGACCG | 61.5 |
| | Reverse | GAGAGTAACGGCCTTTTTGTGA | |
| ***Tmem119*** | Forward | CCTACTCTGTGTCACTCCCG | 62.0 |
| | Reverse | CACGTACTGCCGGAAGAAATC | |
| ***Trem2*** | Forward | CTGGAACCGTCACCATCACTC | 61.5 |
| | Reverse | CGAAACTCGATGACTCCTCGG | |
| ***Gfap*** | Forward | TGGAACAGCAAAACAAGGCG | 60.0 |
| | Reverse | CTGTCTATACGCAGCCAGGT | |
| ***Gja1*** | Forward | GAGAGCCCGAACTCTCCTTTT | 60.0 |
| | Reverse | TGTCTGGGCACCTCTCTTTCAC | |
| ***S100b*** | Forward | TGGTTGCCCTCATTGATGTCT | 61.5 |
| | Reverse | CCCATCCCCATCTTCGTCC | |
| ***Slc1a2*** | Forward | TCCTGGGAGATTGGGTTCAC | 60.0 |
| | Reverse | TATTCTGCCACCCGTTGTCT | |
| ***Il6*** | Forward | CACTTCACAAGTCGGAGGCT | 62.0 |
| | Reverse | CTGCAAGTGCATCATCGTTGT | |
| ***Nos2*** | Forward | CCGAAGCAAACATCACATTCA | 62.0 |
| | Reverse | GGTCTAAAGGCTCCGGGCT | |
| ***Tgfb1*** | Forward | CACTCCCGTGGCTTCTAGTG | 64.0 |
| | Reverse | CTTCGATGCGCTTCCGTTTC | |
| ***Il1b*** | Forward | CCCAAAAGATGAAGGGCTGC | 64.0 |
| | Reverse | AAGGTCCACGGGAAAGACAC | |
| ***Il10*** | Forward | ACCTGGTAGAAGTGATGCCC | 60.0 |
| | Reverse | GCTCCACTGCCTTGCTCTTAT | |
| ***Tlr4*** | Forward | AGATCTGAGCTTCAACCCCTTG | 60.0 |
| | Reverse | AGAGGTGGTGTAAGCCATGC | |
| ***Myd88*** | Forward | AGCCTTTACAGGTGGCCAGAG | 60.0 |
| | Reverse | AAGTTCCGGCGTTTGTCCTAG | |
| ***Cx3cl1*** | Forward | ACGAAATGCGAAATCATGTGC | 61.5 |
| | Reverse | CTGTGTCGTCTCCAGGACAA | |
| ***Tnf*** | Forward | TGTCTACTCCTCAGAGCCCC | 65.0 |
| | Reverse | TGAGTCCTTGATGGTGGTGC | |
| ***Bdnf*** | Forward | TGTGACAGTATTAGCGAGTGGGT | 66.0 |
| | Reverse | TACGATTGGGTAGTTCGGCATT | |
| ***Ccl2*** | Forward | TTA AAA ACC TGG ATC GGA ACC AA | 65.0 |
| | Reverse | GCA TTA GCT TCA GAT TTA CGG GT | |
| ***Cldn1*** | Forward | CTGGGTTTCATCCTGGCTTC | 63.0 |
| | Reverse | TTGATGGGGGTCAAGGGGTC | |
| ***Cldn2*** | Forward | ATACTACCCTTTAGCCCTGACCGAGA | 60.0 |
| | Reverse | CAGTAGGAGCACACATAACAGCTACCAC | |
| ***Cldn4*** | Forward | CGCTACTCTTGCCATTACG | 60.0 |
| | Reverse | ACTCAGCACACCATGACTTG | |
| ***Ocln*** | Forward | AGACTACACGACAGGTGGGG | 60.0 |
| | Reverse | CTGCAGACCTGCATCAAAAT | |
| ***Tjp1*** | Forward | GCAGACTTCTGGAGGTTTCG | 61.0 |
| | Reverse | CTTGCCAACTTTTCTCTGGC | |

### Additional file 1

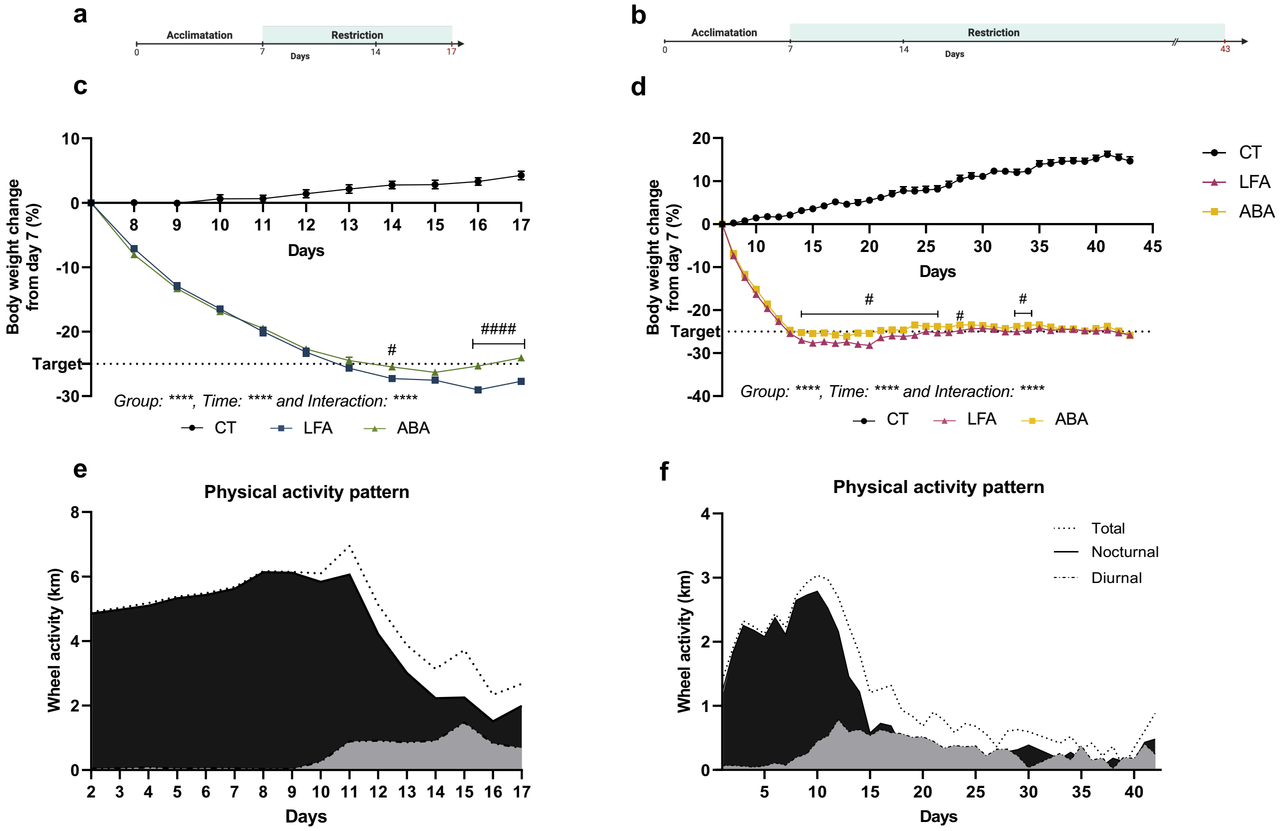


### Additional file 2

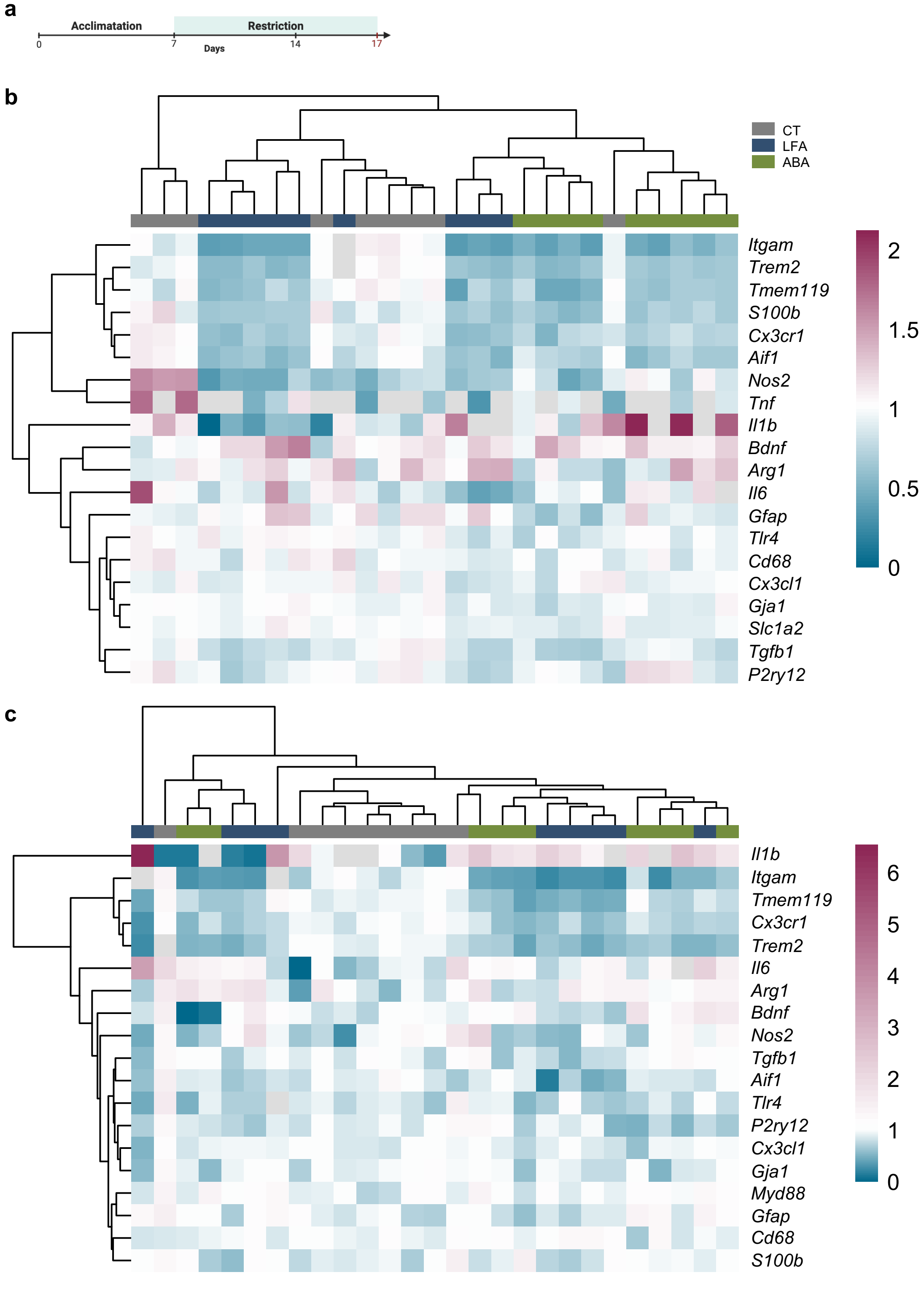


### Additional file 3

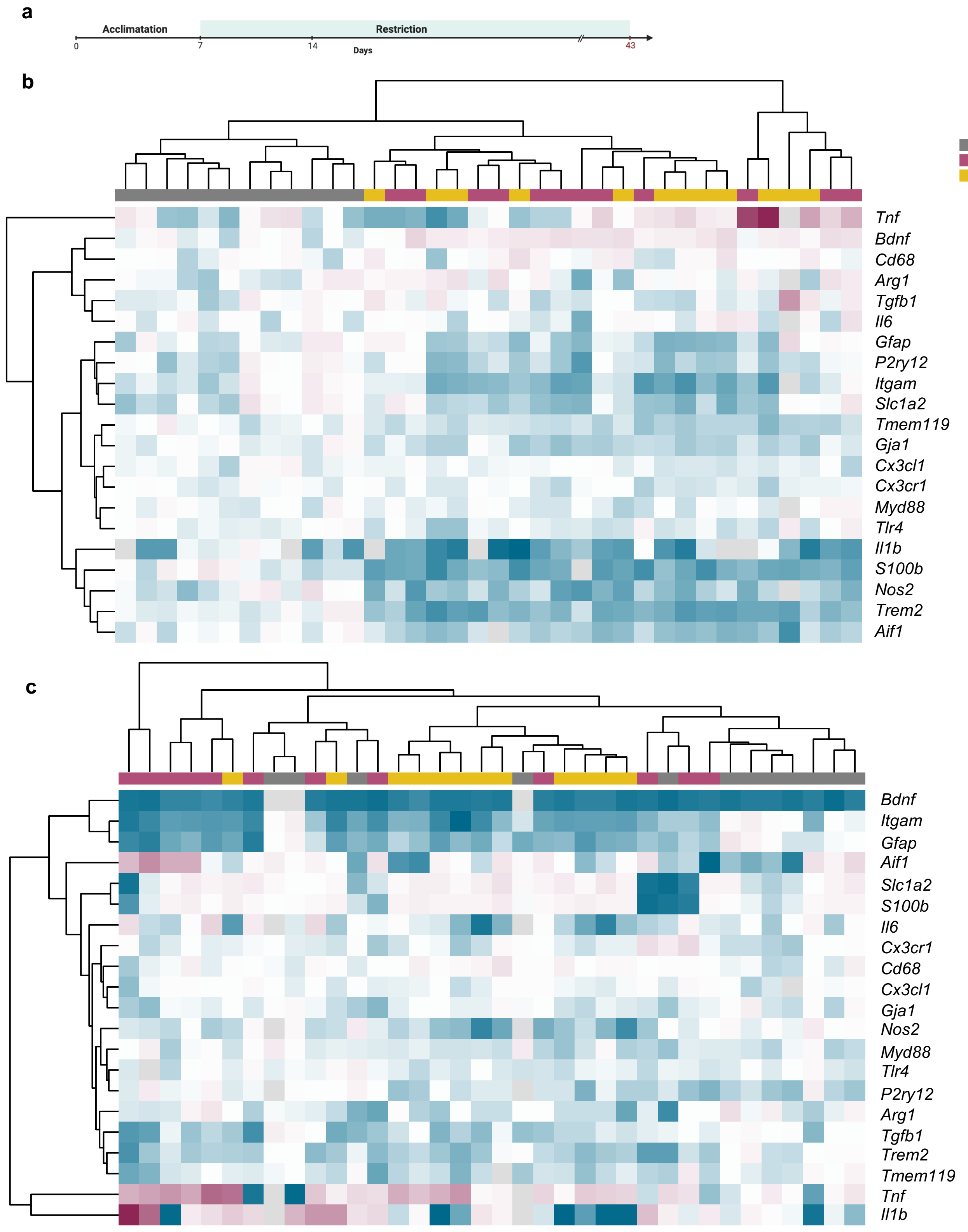


### Additional file 4

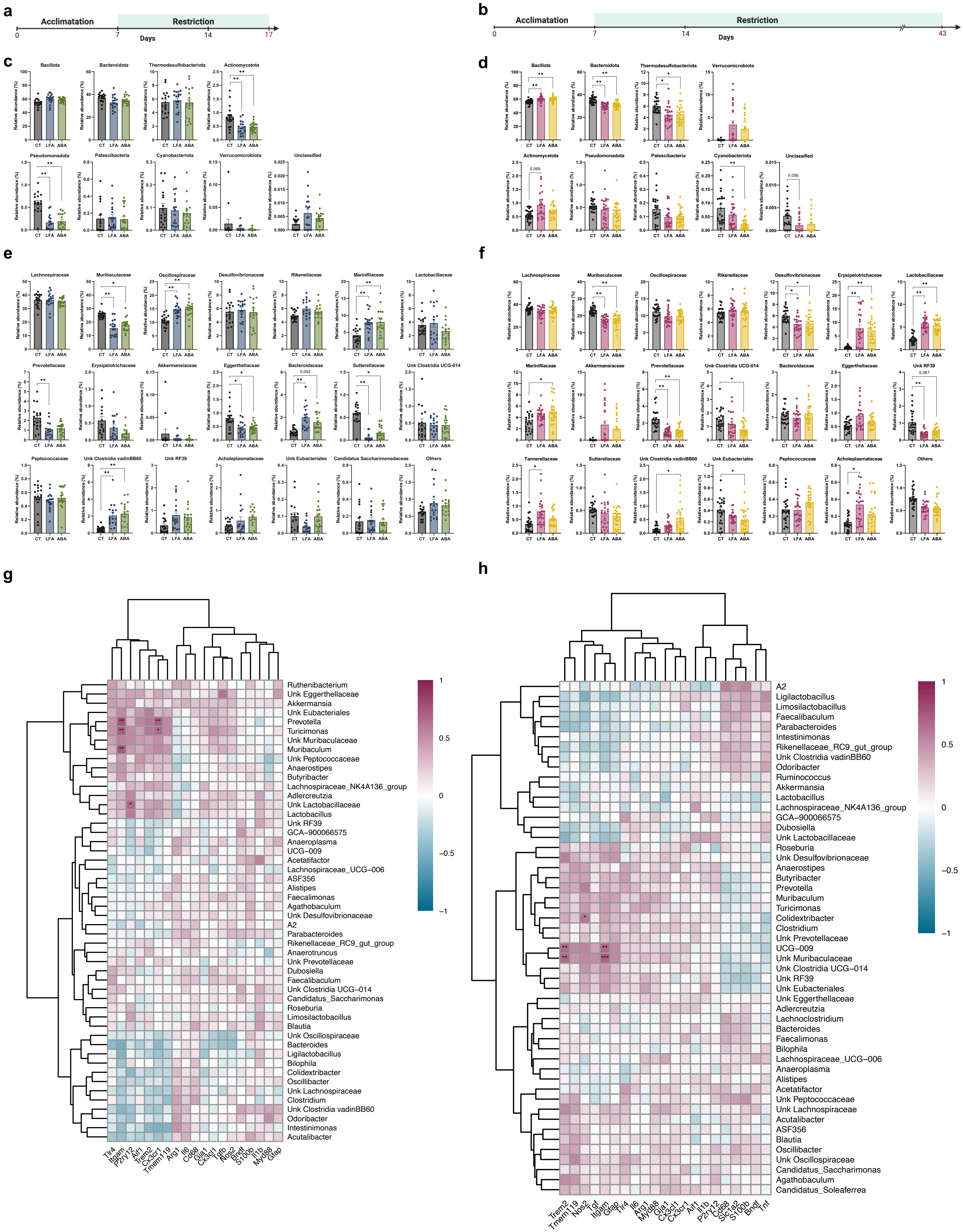


### Additional file 5

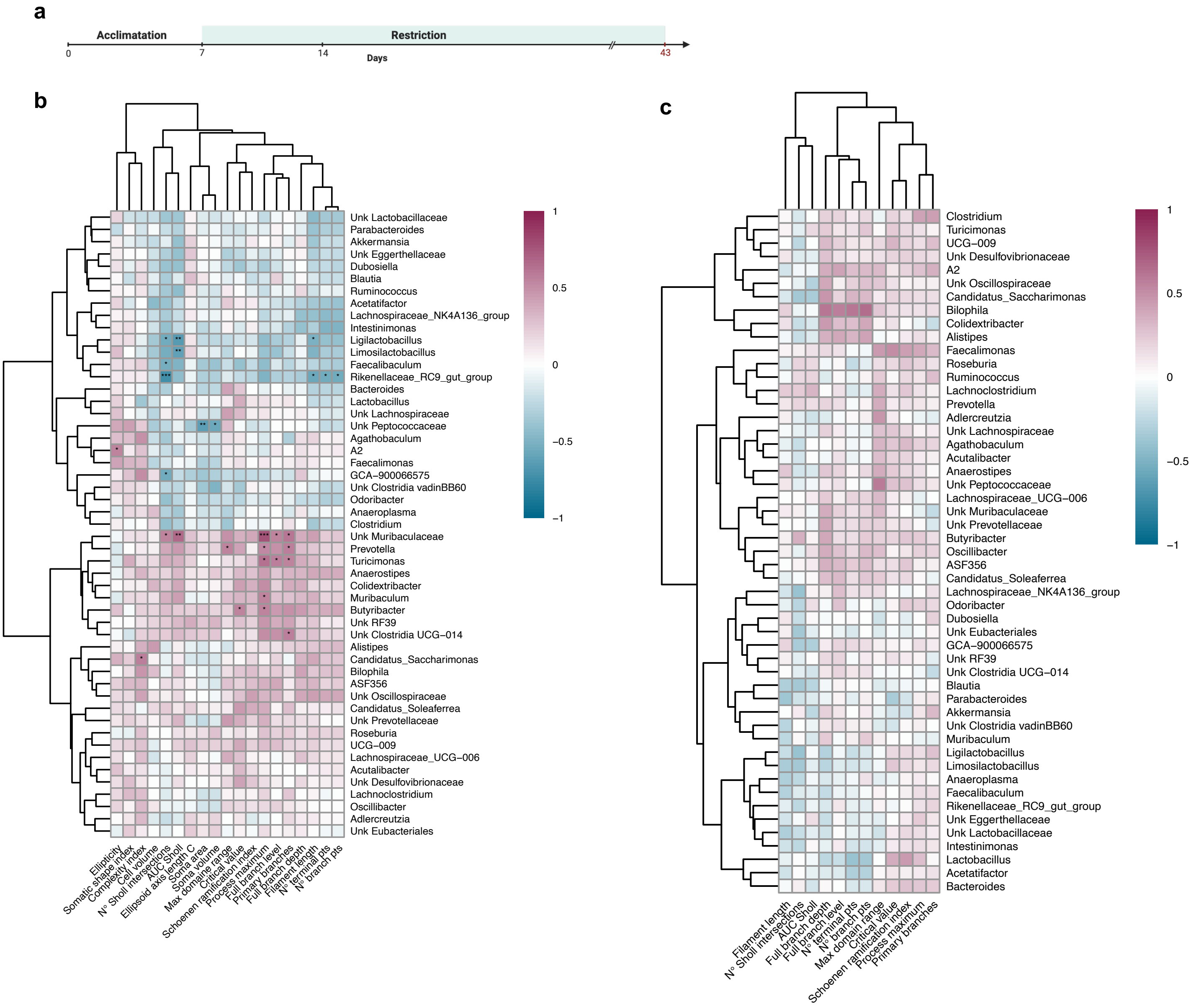


**d**

**Increased**

| Acute | | | | Chronic | | | |
|---|---|---|---|---|---|---|---|
| **LFA and ABA** | *Parent taxa (Phylum, Class, Order)* | | | **LFA and ABA** | *Parent taxa (Phylum, Class, Order)* | | |
| **Marinifilaceae** | *Bacteroidota* | *Bacteroidia* | *Bacteroidales* | **Erysipelotrichaceae** | *Bacillota* | *Bacilli* | *Erysipelotrichales* |
| **Clostridia vadin BB60** | *Bacillota* | *Clostridia* | *Clostridia vadinBB60* | **Lactobacillaceae** | *Bacillota* | *Bacilli* | *Lactobacillales* |
| **Bacteroidaceae** | *Bacteroidota* | *Bacteroidia* | *Bacteroidales* | | | | |
| **Oscillospiraceae** | *Bacillota* | *Clostridia* | *Eubacteriales* | | | | |
| **ABA only** | | | | **ABA only** | | | |
| | | | | **Marinifilaceae** | *Bacteroidota* | *Bacteroidia* | *Bacteroidales* |
| | | | | **Clostridia vadin BB60** | *Bacillota* | *Clostridia* | *Clostridia vadinBB60* |
| **LFA only** | | | | **LFA only** | | | |
| | | | | **Acholeplasmataceae** | *Bacillota* | *Bacilli* | *Acholeplasmatales* |
| | | | | **Tannerellaceae** | *Bacteroidota* | *Bacteroidia* | *Bacteroidales* |

**Decreased**

| Acute | | | | Chronic | | | |
|---|---|---|---|---|---|---|---|
| **LFA and ABA** | *Parent taxa (Phylum, Class, Order)* | | | **LFA and ABA** | *Parent taxa (Phylum, Class, Order)* | | |
| **Muribaculaceae** | *Bacteroidota* | *Bacteroidia* | *Bacteroidales* | **Muribaculaceae** | *Bacteroidota* | *Bacteroidia* | *Bacteroidales* |
| **Eggerthellaceae** | *Actinomycetota* | *Coriobacteriia* | *Coriobacteriales* | **Prevotellaceae** | *Bacteroidota* | *Bacteroidia* | *Bacteroidales* |
| **Sutterellaceae** | *Pseudomonadota* | *Gammaproteobacteria* | *Burkholderiales* | **Unk RF39** | *Bacillota* | *Bacilli* | *RF39* |
| | | | | **Desulfovibrionaceae** | *Thermodesulfobacteriot* | *Desulfovibrionia* | *Desulfovibrionales* |
| **ABA only** | | | | **ABA only** | | | |
| | | | | **Unk Clostridia UCG-014** | *Bacillota* | *Clostridia* | *Clostridia UCG-014* |
| | | | | **Unk Eubacteriales** | *Bacillota* | *Clostridia* | *Eubacteriales* |
| **LFA only** | | | | **LFA only** | | | |
| **Prevotellaceae** | *Bacteroidota* | *Bacteroidia* | *Bacteroidales* | | | | |

### Additional file 6

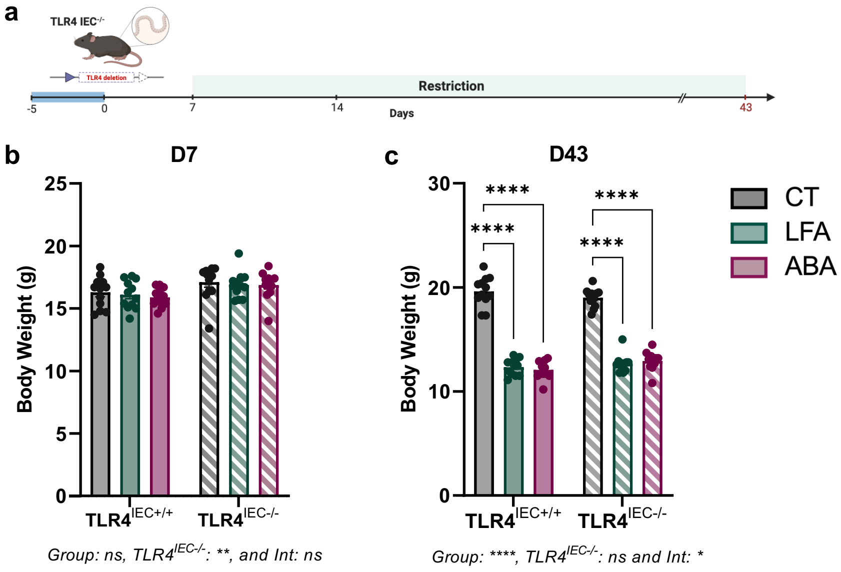


### Additional file 7

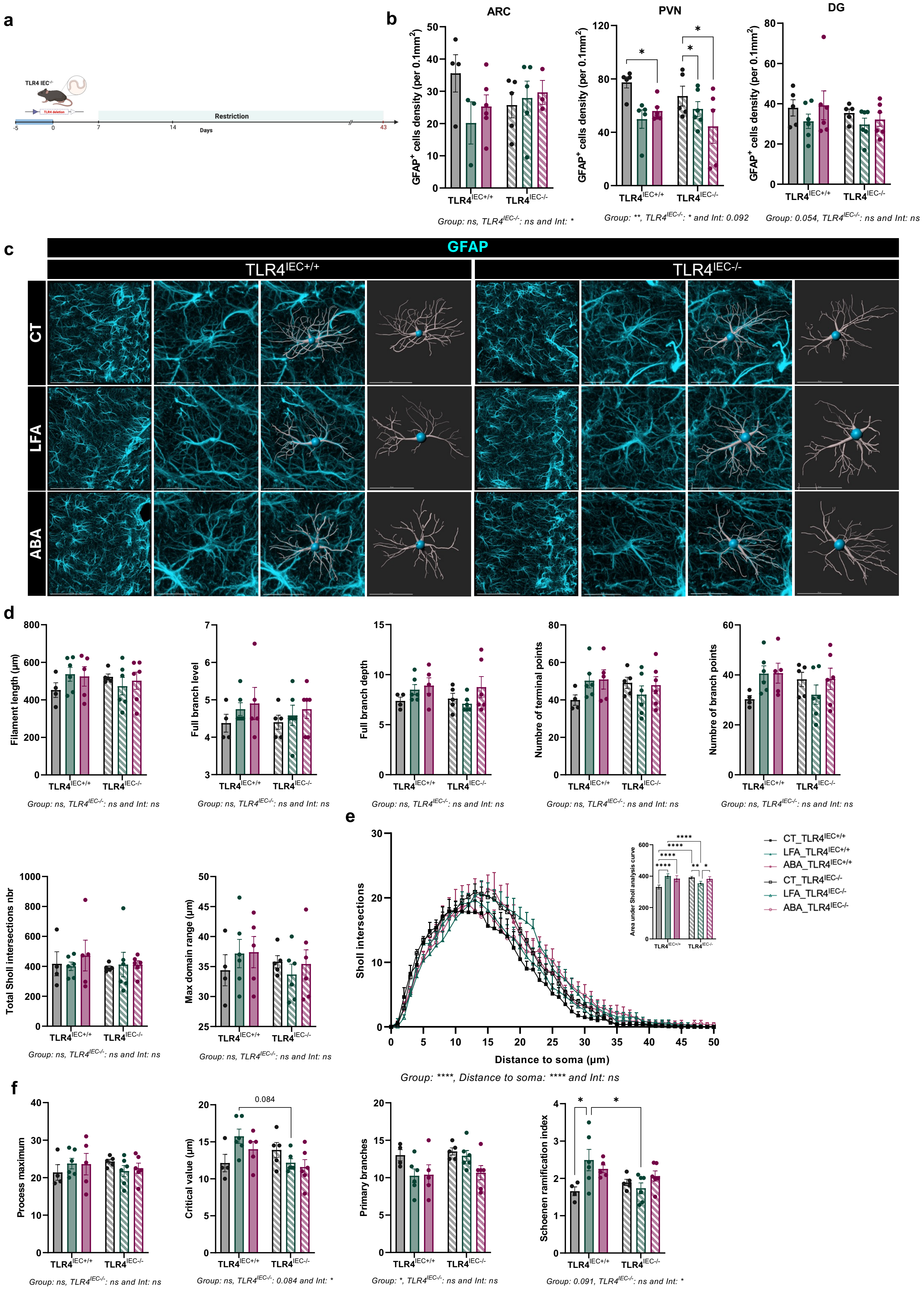


### Additional file 8

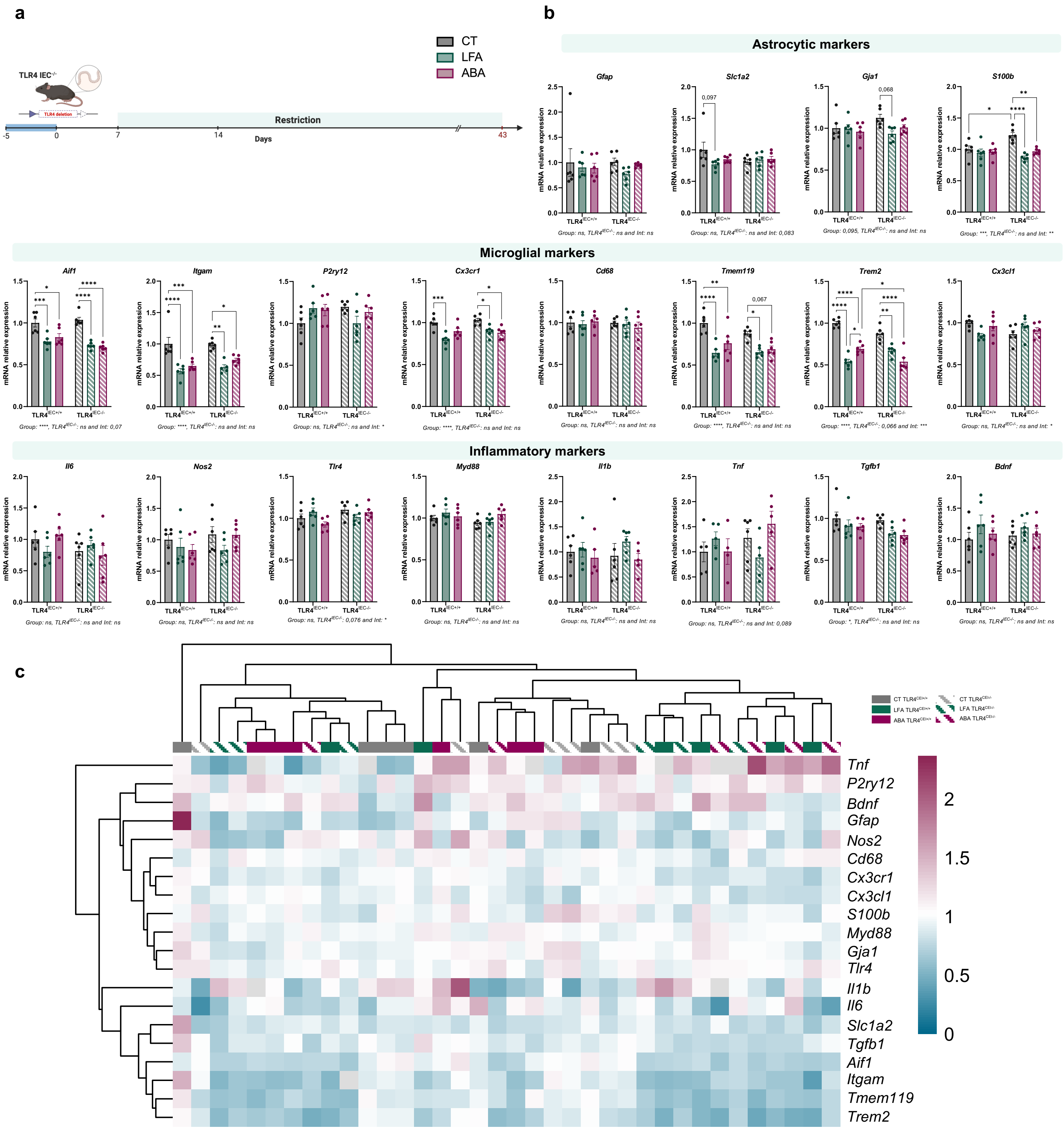

### Additional file 9

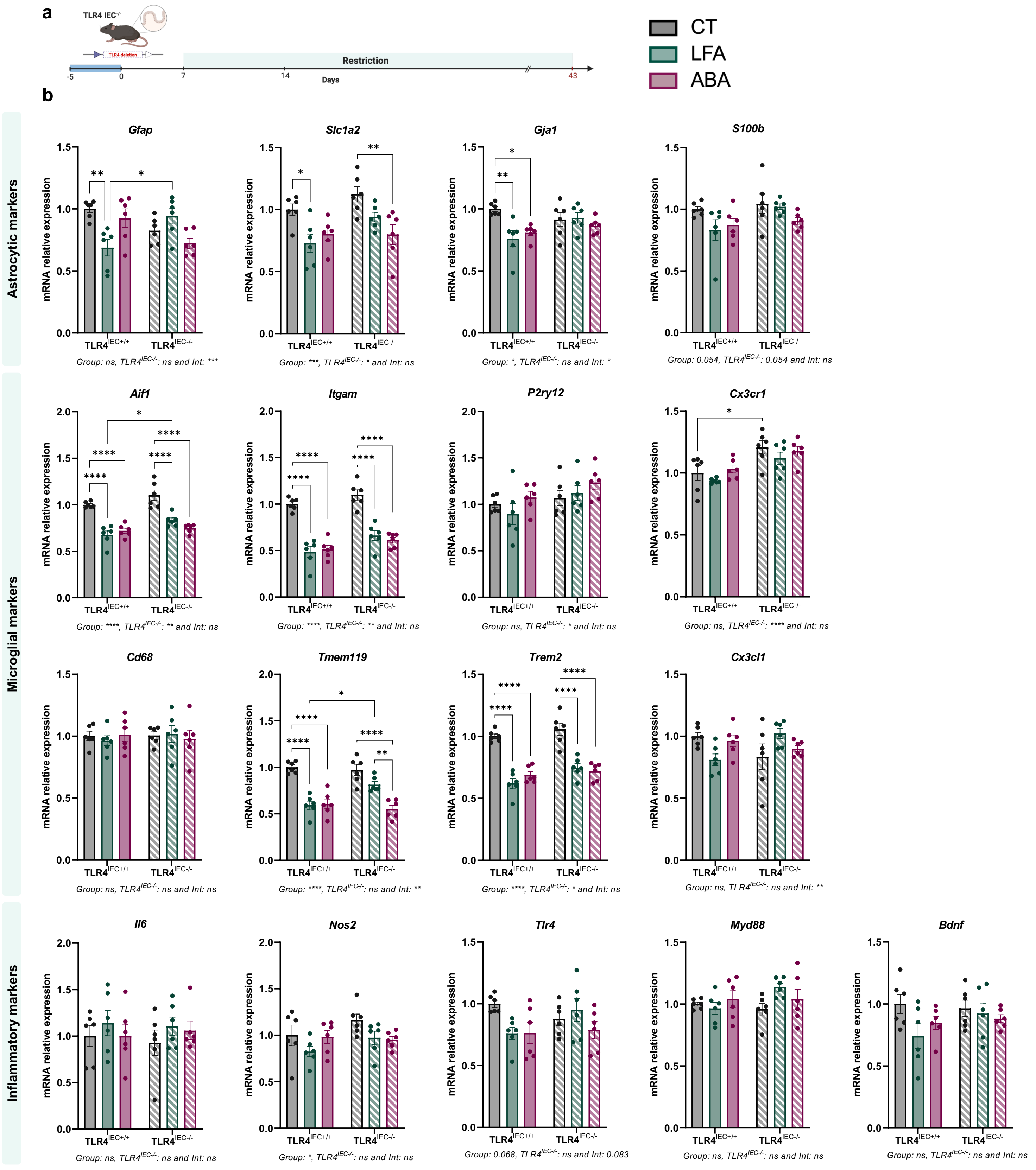


### Additional file 10
